# Conservation and Variations of Bimodal *HoxD* Gene Regulation During Tetrapod Limb Development

**DOI:** 10.1101/372664

**Authors:** Nayuta Yakushiji-Kaminatsui, Lucille Lopez-Delisle, Christopher Chase Bolt, Guillaume Andrey, Leonardo Beccari, Denis Duboule

## Abstract

In all tetrapods examined thus far, the development and patterning of limbs require the activation of gene members of the *HoxD* cluster. In mammals, they are controlled by a complex bimodal regulation, which controls first the proximal patterning, then the distal structure, allowing at the same time the formation of the wrist and ankle articulations. We analyzed the implementation of this regulatory mechanism in chicken, i.e. in an animal where large morphological differences exist between fore-and hindlimbs. We report that while this bimodal regulation is globally conserved between mammals and avian, some important modifications evolved at least between these two model systems, in particular regarding the activity of specific enhancers, the width of the TAD boundary separating the two regulations and the comparison between the forelimb *versus* hindlimb regulatory controls. Some aspects of these regulations seem to be more conserved between chick and bats than with the mouse situation, which may relate to the extent to which forelimbs and hindlimbs of these various animals differ in their functions.

**AUTHOR SUMMARY:** The morphologies of limbs largely vary either amongst tetrapod species, or even between the fore-and hindlimbs of the same animal species. In order to try and evaluate whether variations in the complex regulation of *Hoxd* genes during limb development may contribute to these differences, we compared their transcriptional controls during both fore-and hindlimb buds development in either the mouse, or the chicken embryos. We combined transcriptome analyses with 3D genome conformation, histone modification profiles and mouse genetics and found that the regulatory mechanism underlying *Hoxd* gene expression was highly conserved in all contexts, though with some clear differences. For instance, we observed a variation in the TAD boundary interval between the mouse and the chick, as well as differences in the activity of a conserved enhancer element (*CS93*) situated within the T-DOM regulatory landscape. In contrast to the mouse, the chicken enhancer indeed displayed a stronger activity in fore-than in hindlimb buds, coinciding with the observed striking differences in the mRNA levels. Altogether, differences in both the timing and duration of TAD activities and in the width of their boundary may parallel the important decrease in *Hoxd* gene transcription in chick hindlimb *versus* forelimb buds. These differences may also account for the slightly distinct regulatory strategies implemented by mammals and birds at this locus, potentially leading to substantial morphological variations.

## INTRODUCTION

Tetrapod limbs are organized into three parts bearing skeletal elements; the stylopodium (humerus/femur), the zeugopodium (radius/fibula, ulna/tibia) and the autopodium, the latter including the acropod (phalanges, metacarpals/metatarsals) and the mesopodium (carpals and tarsals)[1]. Limbs can display large variations in their morphologies, either between tetrapod species or within the same species, between fore- and hindlimbs, as a result of their adaptation to different functions and ecological niches. For example, frogs display particular shapes of carpal and tarsal elements, with an elongated proximal tarsal whenever detectable[2], whereas geckos forelimb skeletal elements somehow resemble those of their hindlimbs[3]. Another example of this morphological flexibility are the forelimbs of bats, which do have digits early on similar to those of other mammals but that subsequently elongate to make flight possible[4].

In this context, birds are a fascinating animal group as they evolved forelimbs (wings) and hindlimbs (legs) specialized for flying or for terrestrial locomotion, respectively[5]. Recent studies using comparative genomics approaches either amongst birds or between bats and mice, have revealed that some DNA sequences potentially involved in limb development and which are highly conserved can display differential enhancer activities as compared to their mouse orthologous sequences[6,7]. Furthermore, the analysis of several domestic pigeons displaying variations in foot feathering within the same species, suggested that changes in cis-regulatory elements in the genes encoding forelimb- or hindlimb-specific transcription factors may contribute to a partial transformation from hindlimb to forelimb identity[8]. Taken together, these observations suggest that both the gain of species-specific enhancers, the different activities of the same regulatory sequences and alterations in DNA sequences amongst various species and/or within the same species, contributed to generate these large morphological differences.

In addition to their essential role during axial patterning and organogenesis in vertebrates[9,10], *Hox* genes are required for proper growth and skeletal patterning of tetrapod limbs. In particular, genes belonging to the *HoxA* and *HoxD* clusters for both fore- and hindlimbs, as well as some genes of the *HoxC* cluster during hindlimb development[11,12]. Regarding both *Hoxd* and *Hoxa* genes, recent development in chromatin conformation techniques have made it possible to associate previously defined limb regulatory landscapes to large chromatin interaction domains referred to as TADs (topologically associating domains)[13–15]. Therefore, multiple limb-specific DNA regulatory sequences were identified on either side of the *HoxA* and *HoxD* clusters, grouped either into TADs and smaller sub-megabase large domains (sub-TADs)[16–19].

At the murine *HoxD* locus, two overlapping subsets of genes are controlled by the two adjacent regulatory TADs located on the telomeric side (T-DOM) and the centromeric side (C-DOM) of the cluster[17]. The region of the cluster extending from *Hoxd1* to *Hoxd8* generates constitutive interactions with T-DOM, whereas the 5’ region of the cluster, which includes *Hoxd13* to *Hoxd12,* predominantly contacts C-DOM. Interestingly, the genes from *Hoxd9* to *Hoxd11,* located within the boundary between these two TADs, interact with both T-DOM and C-DOM, yielding their transcripts in both the future zeugopod and acropod domain. In this situation, the expression of *Hoxd1* to *Hoxd11* is driven by contacts with enhancer elements situated in T-DOM in the prospective zeugopod. Then, as the future acropod domain is generated, *Hoxd* genes start to weaken their interactions with T-DOM and *Hoxd9* to *Hoxd11* switch to establish interactions within C-DOM. This switch is partly controlled by HOX13 proteins, which inhibit T-DOM activity while re-enforcing C-DOM-located enhancers function[20]. This bimodal regulatory mechanism allows the production of a domain of low *Hoxd*-expression in which both T-DOM and C-DOM regulations are functionally inert, giving rise to the future mesopodium. While this complex system seems to be globally conserved throughout evolution[21,22], some slight modifications could readily lead to important changes in expression specificities.

In different tetrapod species, the morphological diversifications seen in particular in both the mesopod and the zeugopod between fore-and hindlimbs were suggested to result from heterochronic variations in *Hox* gene expression[2][23]. Both gain of function and loss of function experiments have revealed that either the ectopic expression or the mis-regulation of *Hoxa13* and *Hoxd13* can induce a substantial reduction and malformation of the zeugopod, similar to mesomelic dysplasia conditions in human families (e.g.[24]). This is due to the potential of these particular HOX13 proteins to antagonize the function of other HOX proteins to control and stimulate the ossification of limb skeletal elements[25]. In this view, HOX protein production controlled by the T-DOM stimulate bone growth, whereas C-DOM enhancers upregulate *Hoxd13* to antagonize this property, leading to both smaller bones (phalanges) and the termination of the structure, in a dose-dependent manner[26–30].

In this context, it was recently reported that a bat regulatory sequence located within T-DOM and controlling *Hoxd* genes displays differential enhancer activity in the limbs, when compared to its mouse orthologous sequence[6]. These findings suggest that some variations in limb morphology may be associated with the mechanism of bimodal gene regulation described at the *HoxD* locus. However, this mechanism was reported to be implemented during the development of forelimb buds only and hence it remained unclear as to how much the important differences between fore-and hindlimbs either amongst various species or within the same species, may be related to variations of this mechanism.

To tackle this issue, we used a comparative regulatory approach involving chick and mouse embryonic fore-and hindlimbs, mostly for two reasons. First chicken embryos, unlike mice, display striking differences between the morphologies of their adult wings (forelimbs) and legs (hindlimbs) (Fig. 1A, B, left). Secondly, it was reported that *Hoxd* gene expression domains during chick wing-and leg buds development showed important deviations when compared to their mouse counterparts[23,31]. These features suggested that the bimodal regulatory system at work at the mouse *HoxD* locus may be operating slightly differently during the development of the avian appendicular skeletons.

**Fig. 1.**
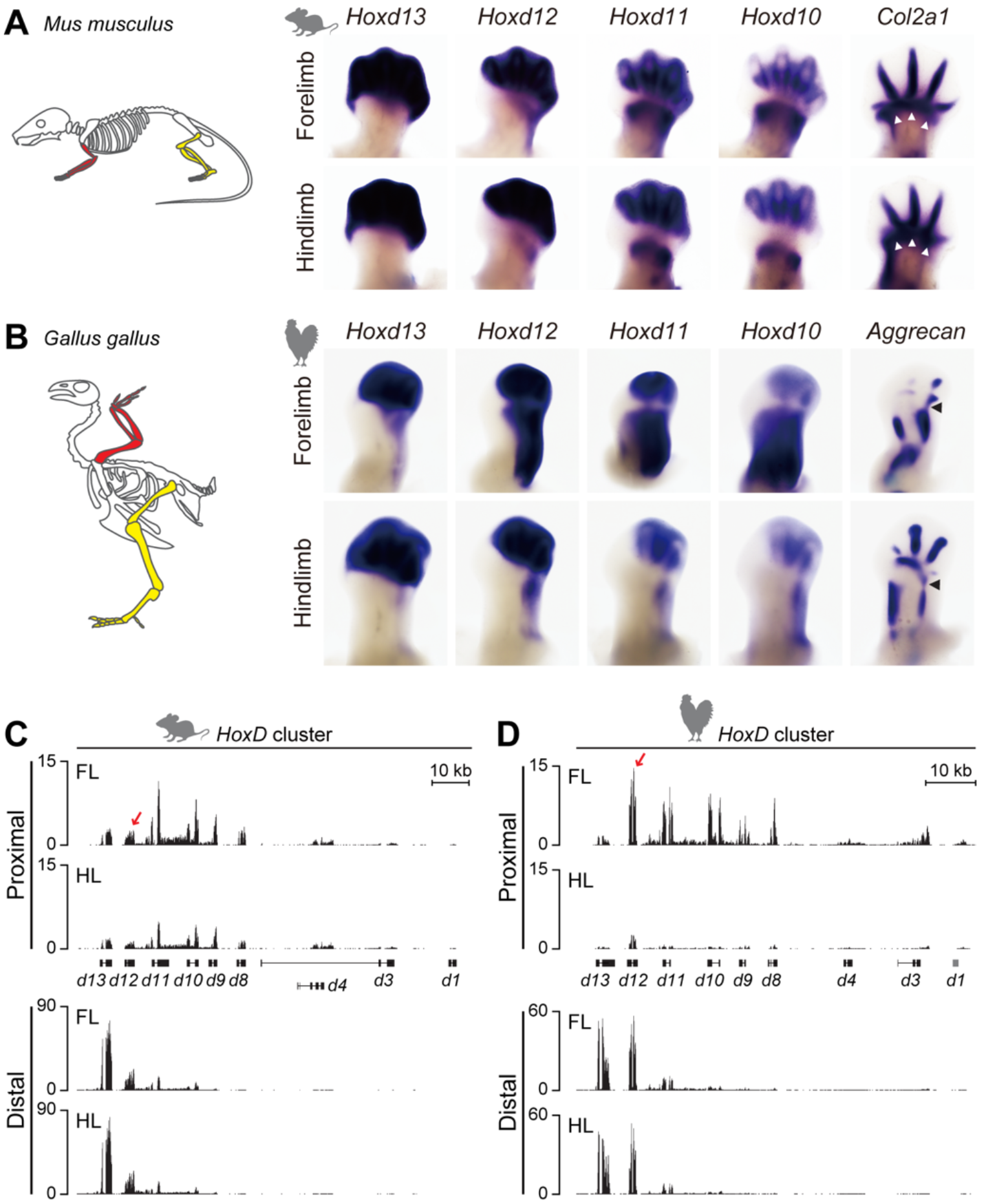
*Hoxd* genes expression in mouse and chick limb buds. (A, B) *In situ* hybridization analysis of E12.5 mouse and HH28 (equivalent to E12.25 to E12.5) chick fore- and hindlimb buds showing the expression of *Hoxd* genes and *Col2a1* or *Aggrecan*, which are markers for chondrocyte differentiation. (A, left) Schemes showing morphologies of forelimb (red) and hindlimb (yellow) in mouse. (A, right) Expression patterns of *Hoxd* genes in forelimb buds are comparable to those in hindlimb buds. The expression domain of *Col2a1* (opened arrowheads) corresponds to the low *Hoxd* expression region leading to the future mesopodium. (B, left) Schemes representing morphologies of forelimb (wing, red) and hindlimb (leg, yellow) buds in chick. (B, right) Expression of *Hoxd* genes in proximal leg is significantly reduced and restricted to the presumptive ulna. (C, D) Transcription profiles of *Hoxd* genes in micro-dissected proximal and distal domains, either from E12.5 mouse (C) or HH30 (equivalent to E13 to E13.5) chick (D) fore- and hindlimb buds. Each right limb in (A, B) is oriented with proximal to the bottom and distal to the top. FL, forelimb; HL, hindlimb. The *Y* axis represents the strand-specific RNA-seq read counts, normalized by the total number of million mapped reads.

Here we combine analyses of transcriptome, 3D genome conformation, histone modification and mouse genetics to show that this bimodal regulatory mechanism is highly conserved in birds. However, in chicken leg buds, the duration of the T-DOM regulation is much reduced, as suggested by dynamic changes in histone modifications, along with a slight difference in chromatin interaction profiles at specific enhancer regions, which accounts for the concurrent reduction in *Hoxd* genes expression in the zeugopod. Finally, by using mutant mouse embryos lacking a large part of T-DOM, we uncovered differences in the effect of T-DOM deletion for *Hoxd* genes between fore-and hindlimbs. Therefore, while these regulatory mechanisms are virtually the same amongst tetrapod species and even within the same species between the fore-and hindlimbs, slight differences are scored, which may partly contribute to the observed morphological differences.

## RESULTS

### Transcription of *Hoxd* genes in mouse and chick limb buds

We first compared the expression patterns of *Hoxd* genes in mouse fore-and hindlimbs at E12.5 (Fig. 1A) with those observed in chick at either HH28 (equivalent to E12.25 to E12.5) (Fig. 1B) or HH30 (equivalent to E13 to E13.5) (Fig. 1D). In mouse fore-and hindlimbs, expression of *Hoxd13* and *Hoxd12* was strong in the prospective acropod region (hereafter termed “distal”), whereas the domains of *Hoxd11* and *Hoxd10* transcripts were split into the distal and zeugopod (hereafter termed “proximal”) regions, except the future mesopod, which was labelled by a *Col2a1*-expressing domain (Fig. 1A, arrowheads). These expression patterns were similar in both fore-and hindlimbs, except for a clearly weaker expression level in the hindlimb proximal domain.

When compared with *Hoxd* gene expression in the mouse counterparts, at least two dramatic differences were confirmed in chick limbs. First, unlike the *Hoxd12* expression pattern observed in murine limbs, the chick *Hoxd12* gene was strongly expressed in chick proximal forelimb (wing) (Fig. 1B). Secondly, the expression of all *Hoxd* genes was significantly reduced in the chick proximal hindlimb (leg) by stage HH28, when compared to chick proximal wing and mouse proximal limbs[23,31]. As a result, the expression domains of the chick *Hoxd12* in wing buds was much like that of *Hoxd11* or *Hoxd10*, in contrast with the mouse where *Hoxd12* was never strongly expressed in the proximal domain. However, the transition between the two *Hoxd-*expressing domains also labelled the future wing mesopod (Fig.1B, arrowheads). Of note, expression of all *Hoxd* genes was weak in proximal leg buds, again in contrast to what was observed during mouse limb bud development (Fig. 1).

To characterize these differences in more detail, we performed RNA-seq analyses by using HH30 limb buds in order to more easily micro-dissect the various domains and thus exclude a potential contamination of the future mesopod from the distal domain. RNA profiles confirmed the differences detected by WISH. First, *Hoxd11* to *Hoxd8* were expressed at lower levels in the mouse proximal hindlimb when compared to forelimb, a situation re-enforced in chick proximal leg where only a weak *Hoxd12* signal was detected (Fig. 1C, D, upper tracks). In contrast, more reads were scored for *Hoxa10* to *Hoxa11* in both mouse and chick proximal hindlimb when compared to forelimb (S1 Fig).

In the distal domains, transcription patterns and profiles from mouse and chick were similar between fore- and hindlimbs, at both *HoxA* and *HoxD* clusters (Fig. 1C, D, lower tracks, S1 Fig). However, the chick profile revealed a consistently higher transcription of *Hoxd12*, which was as strong as *Hoxd13*, whereas the amount of transcripts in the mouse counterpart was much lower than that for *Hoxd13* (Fig. 1C, D, lower tracks). Taken together, these initial results indicated that both the expression status and the transcript domains of *Hoxd* genes displayed significant differences, either amongst species or within the developing fore- and hindlimb buds, in particular in chicken.

### Bimodal regulation in both fore- and hindlimb buds

To determine to what extent these differences could result from variations in the implementation of the bimodal regulatory mechanism, we performed comparative 4C-seq analyses. We used a variety of viewpoints located at comparable positions to reveal potential interactions in both mouse and chicken limb buds. To do this, we cross-annotated *Hoxd* genes regulatory sequences identified in the mouse genome onto the chick genome by using the LiftOver tool in UCSC. These annotations were then used for all following experiments. In both fore- and hindlimbs, interactions were scored between *Hoxd* genes and the regulatory sequences *island III* and *Prox*, as hallmarks of C-DOM transcriptional activity, or with the *CS39* sequence as a proxy for T-DOM activity in either the distal or the proximal region, respectively[17,18]. As seen in mouse forelimbs, *Hoxd11* mainly contacted T-DOM sequences, especially *CS39*, in mouse proximal hindlimb cells, i.e., in cells where T-DOM was fully active and where C-DOM was silent (Fig. 2A, top). In contrast, in mouse distal hindlimb cells, it preferentially interacted with C-DOM sequences such as *island III* and *Prox* (Fig. 2A, bottom). Quantification of contacts indicated 75% of telomeric contacts in proximal forelimb cells and 50% in distal forelimb cells, showing that *Hoxd11* had reallocated 25% of its global interactions toward the C-DOM TAD in distal cells. Likewise, mouse hindlimb cells showed the same interaction profiles, with 70% of telomeric contacts in proximal hindlimb cells and 45% in distal hindlimb cells (Fig. 2A). This comparison indicated that the bimodal regulation is virtually identical between fore- and hindlimbs in mouse.

**Fig. 2.**
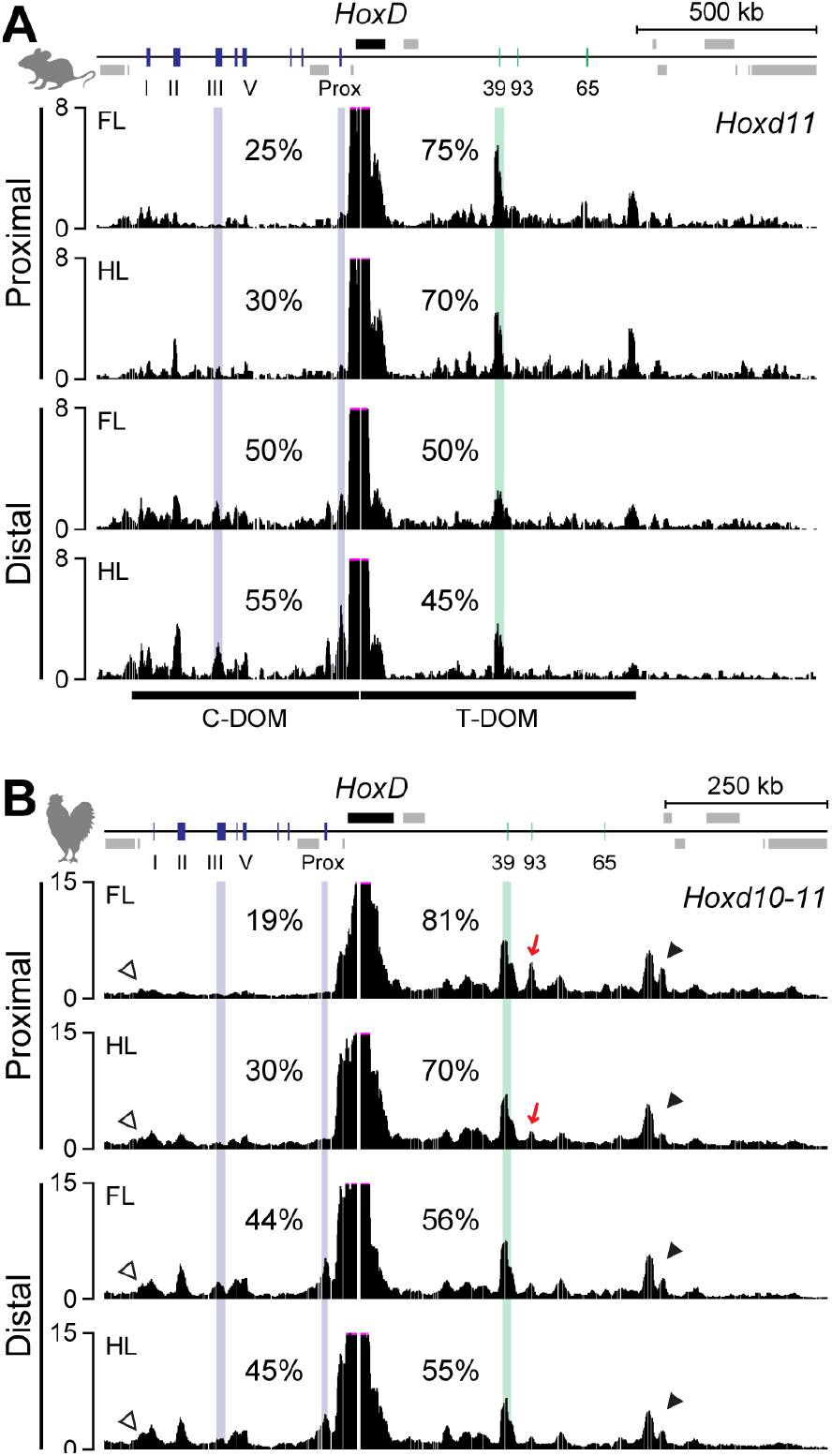
Conserved bimodal TAD regulatory structure at *HoxD* locus in tetrapod. (A, B) 4C-seq tracks showing contacts established by mouse *Hoxd11* and chick *Hoxd10-11* viewpoint in mouse and chick proximal and distal cells from fore- and hindlimb at E12.5 and HH30, respectively. (A) The interactions between *Hoxd11* and around *CS39* were mainly observed in proximal cells, whereas those between *Hoxd11* and *island III* or *Prox*, which are hallmarks of the C-DOM activity, were increased in distal region. (B) Each contact extended to the predicted TAD border located on either side at *HoxD* cluster (C-DOM side, opened arrowheads; T-DOM side, closed arrowheads). In addition to the interactions between *Hoxd10-11* and *CS39*, contacts were also observed with *CS93* in proximal wing bud cells, whereas it decreased in proximal leg bud cells where *Hoxd* expression is strongly reduced (red arrows). FL, forelimb; HL, hindlimb.

We then examined the related interaction patterns in chick wing and leg cells by using a region between *Hoxd11* and *Hoxd10* as a viewpoint (termed ‘*Hoxd10-11’*), i.e. a sequence located as close as possible to the bait used in the mouse experiments. In chick proximal wing cells, *Hoxd10-11* mostly interacted with the *CS39* and *CS93* regions located in T-DOM, as well as with a region near the *Hnrnpa3* gene, where the distal TAD border is observed in the murine locus (Fig. 2B, black arrowhead). These predominant contacts with T-DOM were reduced in chick distal wing cells, as in the mouse, and 25% of contacts were indeed reallocated to C-DOM sequences such as the chicken *island III* and *Prox* counterparts. When compared to chick proximal wing cells, the global interaction with the T-DOM was decreased from 81 to 70% in proximal leg cells. In particular, the interactions between the *Hoxd10-11* bait and the *CS93* sequence in T-DOM were lost in proximal leg cells, which may account for the significant reduction of *Hoxd* expression in chick proximal leg buds (Fig. 2B, red arrows). In contrast, the interaction established by the *Hoxd10-11* bait in chick wing and leg distal cells were comparable, as expected from transcripts analyses, and interactions were observed up to around the *Atp5g3* gene where the border of C-DOM TAD had been mapped in mouse (Fig. 2B, white arrowheads).

### Different enhancer activity of mouse and chick CS93 in fore-*versus* hindlimbs

The mouse *CS93* sequence contains the former *CS9* sequence[17], which was unable to elicit any reporter gene expression in a mouse transgenic context (Fig. 3A). Likewise, a larger murine sequence encompassing *CS9* and referred to as mouse BAR116 (Bat Accelerated Region 116) did not show any enhancer activity in the limbs[6] (Fig. 3A). In contrast, the bat BAR116 sequence was able to drive strong expression in transgenic mouse forelimb buds whereas a weak activity was detected in hindlimb buds, correlating with the different expression levels of *Hoxd10* and *Hoxd11* observed between bat fore- and hindlimb buds[6] (Fig. 3A). This sequence was thus proposed as having evolved a bat-specific function.

**Fig. 3.**
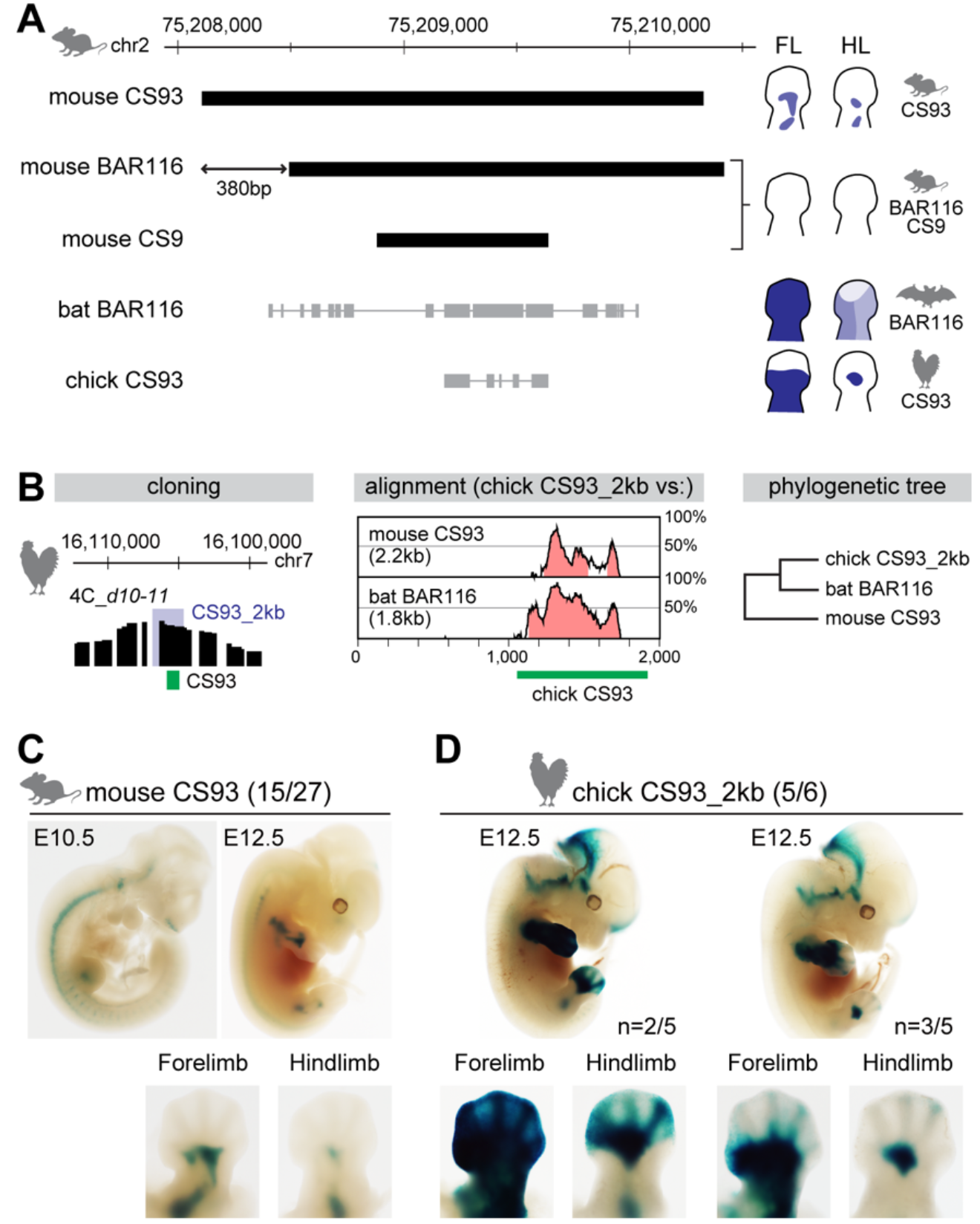
Differential enhancer activities of mouse and chick *CS93* in fore- and hindlimb buds. (A) Genomic coordinates and sequence alignment using either the bat or the chick sequence onto the mouse genome, and schematics of enhancer activities for each of the identified sequences[6,17] and this work. Both mouse BAR116 and mouse *CS9* showed no enhancer activity in limbs[6,17], whereas the bat BAR116 displayed different patterns between mouse fore- and hindlimb[6]. The sequences of both the bat BAR116 (Myoluc2, GL429772: 6,606,808-6,608,652) and the chick *CS93* (galGal5, chr7:16,104,952-16,105,803) were aligned onto the mouse genome. (B) Coordinates of either the chick *CS93* (green rectangle), or the 2kb large region used for enhancer assay (blue domain). The 2kb sequence contains the chick *CS93* region and the region of high interactions with the *Hoxd10* to *Hoxd11* region in proximal wing bud cells at HH30. Conservation plot of mouse *CS93* and bat BAR116 using the 2kb region of chick *CS93* as a reference. The peaks represent a conservation higher than 50%. Pink regions are conserved non-coding sequences. The phylogenetic tree shows the highest conservation of the chick CS93 with the bat BAR116 sequences. (C) Enhancer activities of mouse *CS93* and chick *CS93* in mouse fore- and hindlimb buds from E10.5 to E12.5. The *lacZ* expression pattern (left) showed that mouse *CS93* has an enhancer activity in the proximal region of developing limb buds at E10.5 to E12.5. In contrast to the mouse, the chick *CS93* (right) showed differential enhancer activity between fore- and hindlimb buds at E12.5, as was also reported for the bat BAR116 sequence. The numbers of *lacZ* positive embryos over total transgene integrated are indicated.

Therefore, low expression of *Hoxd* genes in proximal hindlimbs seems to be a common feature of bats and chicken, suggesting that the chick *CS93* sequence may have a limb enhancer activity similar to that reported for the bat BAR116. We examined the enhancer activity of chick *CS93* using a transgenic mouse *lacZ* reporter system and compared it to the activity of full-length mouse *CS93* sequence by using lentivector-mediated transgenesis[32,33]. We initially cloned a 2kb large sequence containing chick *CS93* and those surrounding sequences (Fig. 3B), which showed higher interactions with *Hoxd10* to *Hoxd11* in the 4C profiles obtained from proximal wing cells (Fig. 2B, track 1). We noted that the surrounding sequences are not particularly conserved among species, whereas the *CS93* region of the chick genome is more conserved with the bat than with the mouse counterpart (Fig. 3B). By using the BLAT search tool in UCSC, we also found that most of conserved regions from the bat BAR116 and the chick *CS93* sequences can be aligned onto the mouse *CS9* region, respectively (Fig. 3A).

We assessed their enhancer activities and, unlike for the mouse BAR116, the full-length mouse *CS93* triggered *lacZ* transcription in E10.5 limb buds with an expression localized to the prospective stylopod and zeugopod at E12.5 (Fig. 3C). The staining was weaker in hindlimb than in forelimb buds, possibly due to the delay in limb development[34,35]. Accordingly, the 380bp localized in 5’ of the mouse *CS93* seem to be essential for expression. On the other hand, we found that the 2kb large sequence containing the chick *CS93* displayed limb enhancer activity in mouse limb buds at E12.5 (Fig. 3D). The transgene expression driven by chick *CS93* came under two different patterns. The first one displayed *lacZ* staining throughout the forelimbs (n=2/5), which was similar to the staining observed when the bat BAR116 sequence was assessed in mouse forelimb bud (Figs 3 and 4 in[6]). The second pattern concerned mostly the proximal forelimb buds (n=3/5), as seen when the mouse *CS93* was used (Fig. 3C). In both cases, a weaker expression was observed in hindlimb bud, as in the case of bat BAR116. These results suggest that the downregulation of *Hoxd* genes in chick leg bud involves a weaker activity of- and less interactions with the *CS93* sequence. Taken together with previous research [6], these results indicate that, unlike for the bat BAR116 and the chick *CS93*, the mouse *CS9* is devoid of limb enhancer activity. Accordingly, the 380bp localized in 5’ of the mouse *CS93* seem to be essential for expression.

**Fig. 4.**
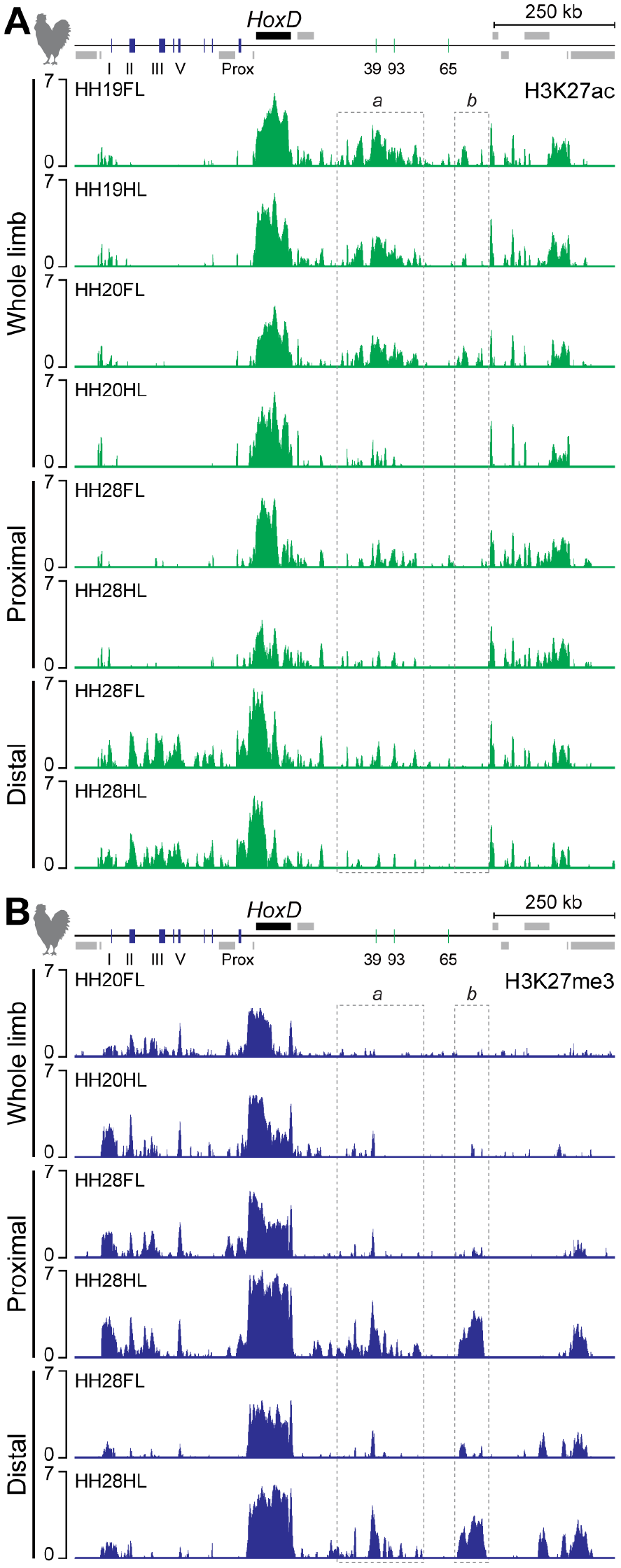
Premature termination of T-DOM activity in chick leg buds. (A, B) Comparison of H3K27ac and H3K27me3 ChIP-seq profiles in either whole, proximal or distal wing- and leg buds at HH19 (equivalent to mouse E9.5), HH20 (equivalent to mouse E10) and HH28. (A) In chick leg bud, enrichment of H3K27ac at region ‘*a’* in T-DOM was initially detected at HH19, whereas it was significantly decreased at HH20. Few H3K27ac marks were scored in region ‘*b’* in hindlimb bud at both HH19 and HH20, as compared with those in wing buds. At HH28, the accumulation of H3K27ac marks was quite low in both the *HoxD* cluster and the T-DOM region in proximal leg, when compared to distal wing cells, whereas the profiles of H3K27ac in the distal region where *Hoxd* genes are strongly expressed were similar between wing and leg buds at HH28. (B) In whole wing- and leg buds at HH20, several C-DOM regions were covered by H3K27me3, whereas T-DOM was either not yet labelled, or started to be enriched around *CS39* in forelimb bud and hindlimb buds, respectively. In proximal leg buds where *Hoxd* expression was reduced, H3K27me3 enrichment was observed at the *HoxD* cluster and over T-DOM, when compared to proximal forelimb buds. Both regions ‘*a’* and ‘*b’* in T-DOM seemed to be specific targets of H3K27me3 enrichment. Enrichment (*Y-axis*) of ChIP is shown as the log_2_ ratio of the normalized number of reads between ChIP and input samples. FL, forelimb; HL, hindlimb.

### Implementation of the regulatory switch between TADs in mouse and chicken

We observed that the differences in *Hoxd12* expression were more substantial between chick fore- and hindlimb buds than in the murine counterparts (Fig. 1C, D). Therefore, we wondered whether the regulatory switch from the T-DOM to the C-DOM regulations would occur differently in the two species. We produced and examined 4C interactions profiles by using *Hoxd12* itself as a bait. Similar to the profiles obtained with the *Hoxd10-11* bait, we observed weaker interactions between *Hoxd12* and both the *CS39* and *CS93* regions in T-DOM in chick proximal leg cells than in proximal wing cells (S2A Fig, top). The profiles with the *Hoxd10-11* bait showed strong and stable interactions with T-DOM, when compared with C-DOM, in both proximal and distal limbs (Fig. 2B). We also found that *Hoxd12* mainly contacted T-DOM in both chick proximal wing and leg cells, whereas it established more interactions with C-DOM in both chick distal cells (S2A Fig, top and bottom).

The murine *Hoxd9* to *Hoxd11* genes, but not *Hoxd12*, are located within the TAD boundary and interact both with T-DOM and with C-DOM. In contrast, in chicken limb buds, *Hoxd12* was able to switch contacts from T-DOM to C-DOM, suggesting that the TAD boundary in chick could be located between *Hoxd12* and *Hoxd13* (see also S2B and S2C Fig) thus suggesting that the exact genomic interval where the switch in contacts occurs between the genes and the TADs is displaced towards a more 5’ position within the chicken gene cluster, when compared to murine limb buds where this switch region was localized around the *Hoxd11* locus[17,19]. This same switch was observed in both chick wing- and leg bud cells, regardless of the various expression levels of *Hoxd* genes in the proximal region, revealing that the switch between TADs is independent of *Hoxd* expression itself in proximal cells.

These results showed that the bimodal regulatory mechanism and the sequential transition from the proximal to the distal global controls are implemented during chick limb development similarly to what was described in mice. Therefore, the differences in gene expression observed both between mice and chicken and between chick wing- and leg buds cannot be solely explained by visible variations in the respective interaction profiles. Instead, they ought to involve the distinct use of enhancers (or group thereof-) within an otherwise globally conserved chromatin architecture.

### Premature termination of the proximal TAD activity in chick leg buds

Since the chromatin architecture at the *HoxD* locus is seemingly comparable between mouse and chicken in both fore- and hindlimb buds, we looked for what may cause the drastic reduction of *Hoxd* expression observed in chick proximal leg. Within the T-DOM TAD structure, the interaction profiles obtained from chick leg proximal cells showed reduced contacts between *Hoxd* promoters and enhancers in T-DOM (Fig. 2 and S2 Fig). We complemented these observations by assessing the functional state of T-DOM sequences by comparing particular histone modifications profiles between chick wing- and leg buds, at several developmental stages (Fig. 4). We looked at the acetylation of H3K27 (H3K27ac), a modification associated with transcriptional and enhancer activity and at the trimethylation of the same residue (H3K27me3), a mark associated with Polycomb-dependent silencing[36]. In both wing- and leg buds at stage HH19, a stage which corresponds to ca. E9.5 in mouse, enrichments of H3K27ac were detected over both T-DOM and the *HoxD* cluster itself, showing that the activation of *Hoxd* genes by the T-DOM enhancers had been properly initiated in leg buds (Fig. 4A, tracks 1 and 2). Of note, higher levels of this mark were scored over the *Hoxd11* to *Hoxd13* region in leg-than in wing buds (S3A Fig).

At stage HH20 (approximatively E10 in mouse), the H3K27ac enrichment in T-DOM was still substantial in wing buds. In marked contrast, however, this level appeared dramatically reduced in leg buds (Fig. 4A, tracks 3 and 4, region *a*), thus coinciding with low gene expression (S3A Fig). The accumulation of H3K27ac observed near the TAD border was specific for the early wing-bud (Fig. 4A, tracks 1 and 3, region *b*). Furthermore, H3K27ac signals over C-DOM were not yet observed at these stages (except around the island I region), in agreement with the fact that the regulatory switch had not yet occurred. At a later stage (HH28, i.e. approximately E12.5 in mouse), enrichment of H3K27ac within the *HoxD* cluster was significantly lost in proximal leg bud cells where *Hoxd* expression was weak (Fig. 4A, track 6, S3A Fig). In contrast, H3K27ac accumulation over the T-DOM in proximal wing bud cells remained, yet it started to slowly decrease as observed in mouse proximal forelimb at E12.5 (Fig. 4A, track 5). At the same time, H3K27ac was finally enriched over both C-DOM and the *HoxD* cluster, in both wing- and leg distal cells, as scored in mouse distal forelimb buds (Fig. 4A, tracks 7 and 8, S3A Fig)[17,20]. These various profiles showed that in chick leg bud cells, the functional switch between T-DOM and C-DOM had occurred normally, except that after its initial onset, T-DOM activity was terminated much more rapidly than in the wing bud, followed by a decrease in accumulation of H3K27ac at the target *HoxD* cluster itself.

We complemented these observations by analyzing H3K27me3 marks, which antagonize the acetylation of the same residue[36]. At stage HH20, no H3K27me3 signal was detected over T-DOM in both wing- and leg buds, except around *CS39* in leg buds (Fig. 4B, tracks 1 and 2, region ‘*a’*), in agreement with the H3K27ac profiles (compared with Fig. 4A, tracks 3 and 4). In contrast, strong levels of H3K27me3 enrichment were observed over the C-DOM regions, where H3K27ac peaks were not detected (Fig. 4B, tracks 1 and 2), suggesting that the activation of *Hoxd* genes by C-DOM regulation had not yet occurred at this stage.

At the *HoxD* cluster itself, stronger levels of H3K27me3 enrichment were clearly detected in leg buds, as compared with wing buds, from the pseudo-*Hoxd1* gene to *Hoxd8*, a DNA interval controlled by T-DOM regulation (S3B Fig). At later stages, H3K27me3 marks were observed over C-DOM in proximal wing bud cells, where C-DOM is inactive, whereas the levels of H3K27me3 marks over T-DOM, in both proximal and distal wing bud cells, were comparable to those seen in the H3K27ac profiles (Fig. 4B, tracks 3, 5). We had previously suggested that the binding of HOX13 proteins to sequences located in T-DOM could cooperate with Polycomb complexes to terminate T-DOM regulatory activity[20].

Altogether, the distribution of both H3K27ac and H3K27me3 marks in chicken limb buds were in agreement with the observed expression status of *Hoxd* genes profiles. A major difference was scored, however, when compared to their mouse counterparts. In proximal leg bud cells at HH28 indeed, where *Hoxd* gene expression is quite weak (Fig. 4B, track 4; regions ‘*a*’ and ‘*b*’), T-DOM and the *HoxD* cluster were heavily decorated with H3K27me3 marks, in addition to the C-DOM TAD. The latter profile resembled that obtained from distal leg bud cells at the same stage, i.e. cells where T-DOM is inactive and completely shut down (Fig. 4B, track 6; regions ‘*a*’ and ‘*b*’), suggesting that T-DOM was not operational in proximal leg cells, unlike in the mouse forelimb proximal situation[17,20].

We also examined the distribution of both H3K27ac and H3K27me3 marks over the *HoxA* cluster and its limb regulatory landscape that maps within a sub-TAD adjacent to *Hoxa13* [16](S4 Fig). H3K27ac distribution suggested an earlier activation of *Hoxa13* in leg-than in wing buds at HH19 and HH20 (S4 Fig), which corroborated the transcriptomes established at HH20 (S4C Fig). At HH28 however, the profiles of H3K27ac and H3K27me3 marks over the *HoxA* cluster and within the flanking regions were comparable between wing- and leg bud cells (S4 Fig) and did not show any of the differences reported for the *HoxD* cluster and their regulatory landscapes. H3K27ac profiles were similar over the *HoxA* limb enhancers in wing- and leg tissues at all stages, except perhaps for a slight decrease in H3K27ac in leg bud at HH20 (S4A Fig).

### Chromatin conformations of the chicken *HoxD* cluster in wing- and leg buds

Gene expression usually occurs concomitantly with enhancer-promoter contacts[37–39]. Because of the dramatic difference in T-DOM activity observed in chick leg-*versus* wing buds at stage HH20 (Fig. 4A, tracks 3 and 4, region ‘*a*’), we looked for potential parallel differences in chromatin contacts by performing high-resolution capture Hi-C analyses (CHi-C)[40,41] using wing- and leg buds at HH20. Since such a global chromatin assessment had not been done during chick development, it also allowed us to compare it with mouse counterpart cells and see to what extent these complex regulatory systems were conserved in distinct groups of tetrapods.

The CHi-C profiles of chicken cells confirmed that the *HoxD* cluster is positioned at the boundary between two TADs, similar to what was proposed in mouse limb bud tissues[17,19] and the two sub-TADs seen in the murine T-DOM were detected as well (S5A and S5B Fig). To position the boundary between the two TADs, we applied the TopDom algorithm[42], which determined the border around the *Hoxd13* locus in both wing- and leg bud cells at HH20 (Fig. 5A). This extended the conclusion reached after the 4C analyses, i.e. that the TAD boundary region in chick was displaced towards the 5’ part of the gene cluster when compared to mouse limb bud cells[19].

**Fig. 5.**
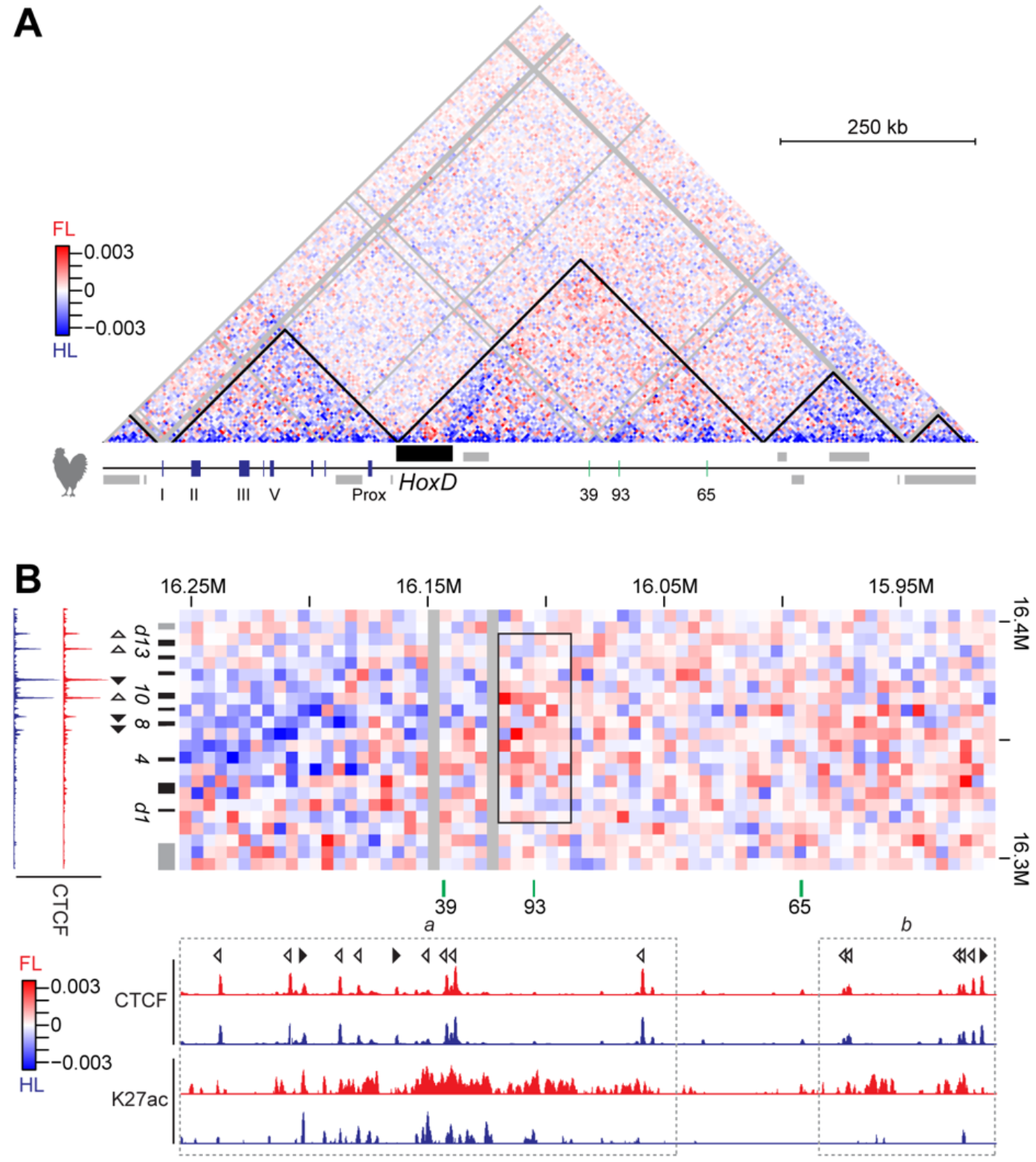
Subtraction of CHi-C matrices between wing- and leg buds with plotted TADs. (A) The black lines demarcate the TADs in wing- and leg buds as identified using the TopDom algorithm. Subtraction are those of CHi-C matrices shown in S5A and S5B Fig, with wing bud cells in red and leg bud cells in blue. (B) Subtraction of the CHi-C matrices shown in S5A and S5B Fig at regions between the *HoxD* cluster and the area ranging from region *‘a’* to region *‘b’* within T-DOM, with a 5kb resolution. A significant decrease in contacts was detected between the *HoxD* cluster and the *CS93* region, which corresponded to the reduction in H3K27ac levels (black rectangle). The increased contacts involving the *HoxD* cluster shown in red and blue represent wing- and leg buds cells, respectively. ChIP-seq profiles of CTCF and H3K27ac from wing- and leg buds at HH20 are shown in red and blue, respectively. Opened and closed arrowheads indicate the orientation of the CTCF motives. Enrichments (*Y-*axis) of CTCF and H3K27ac ChIP are shown at the normalized 1x sequencing depth or the log_2_ ratio of the normalized number of reads between ChIP and input samples, respectively. FL, forelimb; HL, hindlimb.

In mouse limb cells, this TAD boundary falls within a region where multiple CTCF sites are occupied[43–45]. CTCF is an architectural protein that both helps defining constitutive domains of interaction and triggers enhancer-promoter contacts[46]. We thus examined the presence of bound CTCF at the chick *HoxD* locus and surrounding TADs and found that the profiles were comparable between wing- and leg buds at HH20 (S5 Fig). As for the mouse *HoxD* cluster[19], the orientations of the CTCF motives located on either sides of the TAD boundary were facing those sites found in their flanking TADs, suggesting the possibility for long-range loops to be established (S5C Fig, top)(e.g.[47]). Noteworthy, the orientation of the CTCF motives were conserved between mouse and chick. However, we found less bound CTCF in the chicken *HoxD* cluster than in the mouse counterpart, which could affect the strength and/or stability of the TAD boundary in chick (S5C Fig, bottom).

Using CHi-C at 5kb resolution, the distribution of contacts was relatively similar between wing- and leg bud cell populations, despite the reduced level of H3K27ac in the T-DOM and near the TAD border in leg bud cells described above. The contacts between the *HoxD* cluster (galGal5, chr7:16,315,000-16,395,000) and the T-DOM region ‘*a’* (galGal5, chr7: 16,045,000-16,255,000) were indeed comparable between wing- and leg bud cells (Wilcoxon sum test, W=222,690, p-value=0.663) (Fig. 5B). However, when the comparison was further limited to a 30kb large region including *CS93* (galGal5, chr7:16,090,000-16,120,000) rather than to the entire region ‘*a*’, the contact intensity in leg bud cells was clearly below that scored in wing bud cells (W=5,495, p-value=0.0213). These reduced contacts between *Hoxd* genes and the surroundings of region *CS93* confirmed the observation made by looking at the 4C profiles obtained using the later stage (HH30) (Fig. 2).

The fact that bound CTCF was not detected around the *CS93* region suggests that CTCF-independent variations in enhancer-promoter interactions may participate to the important decrease in *Hoxd* gene expression levels in leg bud cells. The decreased contacts seen in leg bud cells were also scored between the *HoxD* cluster and around the T-DOM border region *‘b’* (galGal5, chr7: 15,910,000-15,985,000; W=32,746, p-value=0.009415). The decrease in contacts between the *HoxD* cluster and both the *CS93* and TAD border regions in leg bud cells was somehow compensated for by an increase of interactions between the *HoxD* cluster and the 5’ region of *CS39* where several bound CTCF sites were observed. The 4C profiles obtained when using the *Hoxd10-11* bait showed interactions with the 5’ region of *CS39* in both chick proximal and distal cells, suggesting that these stable contacts are associated with CTCF, as described in mouse developing limb buds[48].

### Regulation of T-DOM by HOXA13

We asked what may cause both the robust reduction in H3K27ac marks in chicken T-DOM and the decrease in contacts between *Hoxd* genes and the *CS93* region in leg bud cells at HH20. As it was reported that HOX13 proteins bind T-DOM-located sequences concomitantly to its inactivation and that the absence of these proteins prevented T-DOM to terminate operating[20,49], we re-assessed the expression dynamics of *Hoxa13*. We found it expressed earlier in chick leg bud than in wing buds[31](S4C Fig), suggesting that the exact timing of *Hoxa13* transcriptional activation may fix the duration of T-DOM activity during limb development.

We examined this possibility by performing time-course *in situ* hybridization and RT-qPCR experiments, using chick and mouse entire fore- and hindlimb buds from HH20 to HH22 and E10.5 to E10.75, respectively (Fig. 6). While these developmental stages are not strictly equivalent between chick and mouse[50], they were selected because the size difference between chick wing- and leg buds is not yet too large between HH20 and HH22[51]. Also, *Hoxa13* starts to be expressed in mouse forelimb buds at around E10.5[52]. While the onset of *Hoxa13* expression was detected in chick wing bud at HH22, *Hoxa13* transcripts were already well present in chick leg bud at HH20-21 (Fig. 6A, left). Also, the expression level of this gene in leg buds was markedly stronger than in wing buds (Fig. 6A, right). *Hoxa11* expression was also higher in chick leg buds than in forelimb buds, as was also observed in the RNA-seq dataset (S4C and S6A Fig), suggesting that the entire chicken *HoxA* cluster was activated in leg buds well before it was switched on in wing buds. This was nevertheless not a general phenomenon for *Hox* genes, for the expression of *Hoxd13* was comparable between wing- and leg buds (Fig. 6A, bottom and S6B, S6C Fig, left).

**Fig. 6.**
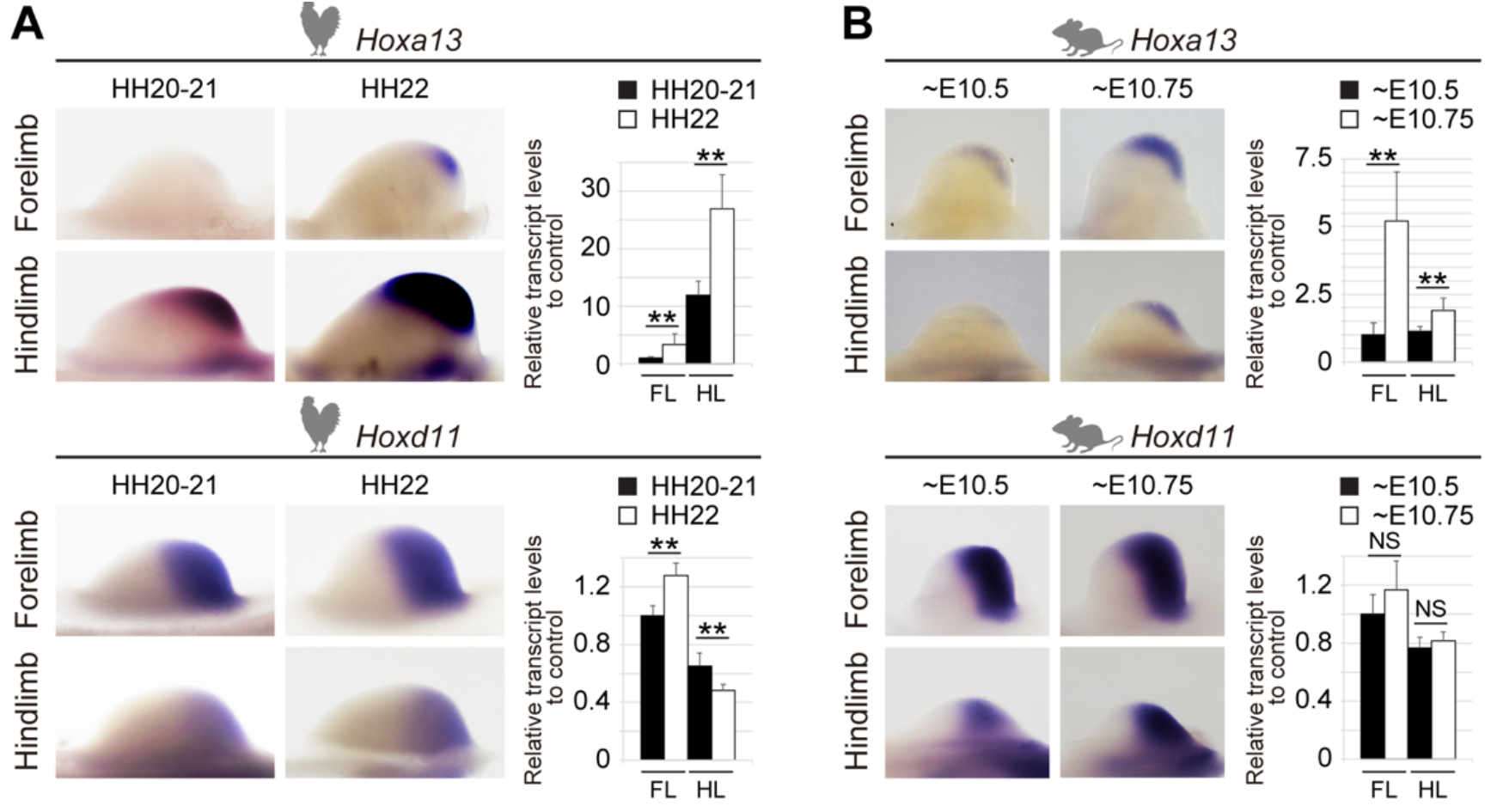
Leg bud-specific early activation of the chick *Hoxa13*. (A) Expression patterns and levels of *Hoxa13* and *Hoxd11* in chick wing- and leg buds from HH20 to HH22. Stronger expression of *Hoxa13* was observed in chick leg bud as compared to that in wing bud (top). Expression level of *Hoxd11* increased in wing bud as development proceeded, yet it decreased in leg bud (bottom). Expression levels were normalized to *Gapdh* and depicted as fold change relative to wing bud at HH20-21. Error bars indicate standard deviation of three biological replicates. **P<0.01, Student’s *t*-test. (B) Expression of *Hoxa13* and *Hoxd11* in mouse fore- and hindlimb buds, from E10.5 to E10.75. Both genes were increasingly transcribed in fore- and hindlimb buds as development proceeded. Expression levels were normalized to *Gapdh* and depicted as fold change relative to forelimb buds at E10.5. Error bars indicate standard deviation of two or four biological replicates. **P<0.01, NS, P>0.05, Student’s *t*-test.

In the mouse, the development of the forelimb bud precedes that of hindlimb buds by about half a day. In contrast, the initiation of both wing- and leg bud in chicken is almost concomitant and the growth of the leg bud exceeds that of the wing bud[34,51]. However, even when considering these developmental differences, the dramatic variations we scored between both the timing of *Hoxa13* activation and its transcript levels between the chick wing- and leg buds were different from the situation observed in murine fore- and hindlimb bud (Fig. 6B, top). Also, expression of other *Hoxd* genes increased in both mouse fore- and hindlimb buds, as development proceeded (Fig. 6B bottom, S6B and S6C Fig. right), unlike in chicken leg buds. Therefore, an inverse correlation exists between the activation of *Hoxa13* on the one hand, and the downregulation of *Hoxd* genes such as *Hoxd11* in chick leg bud on the other hand. This correlation was observed neither in chick wing bud, nor in mouse limb buds, supporting the idea that an early activation of *Hoxa13* induces a premature termination of T-DOM activity in chick leg bud.

We looked whether the profiles of H3K27ac ChIP-seq and Hi-C data obtained from chick limb tissues and covering the *HoxA* cluster would reveal traces of this early and strong activation of *Hoxa13* seen in chick leg buds at HH20 (S4A and S6D Fig). While this activation was consistent with enriched H3K27ac marks over the *HoxA* cluster itself, it was not fully consistent with the distribution of chromatin marks over those enhancers previously described to regulate *Hoxa13* in developing mouse limbs[16](S4A Fig). Hi-C heat maps, however, showed increased contacts between the 5’ region of the *HoxA* cluster and its limb enhancer regions in leg bud at HH20, when compared to wing buds, suggesting this may contribute to an early activation of *Hoxa* genes in leg buds (S6D Fig).

### Different impacts of T-DOM upon mouse fore- and hindlimb buds developments

The importance of the T-DOM TAD for mouse proximal limb development was initially assessed in forelimbs exclusively. The fact that birds displayed this striking difference in T-DOM-dependent regulations in wing and leg buds suggested that tetrapod fore- and hindlimbs may use the T-DOM enhancers in a slightly different manner. We investigated this possibility by looking at the effect of a full deletion of T-DOM upon *Hoxd* gene regulations in both murine fore- and hindlimb buds, in order to identify a potential difference in the genetic requirements for T-DOM regulations between the two contexts. We analyzed mouse limb buds carrying the *Hox^DDel(attp-SB3)^* allele, where an approximatively 1Mb large region including the entire T-DOM as well as its distal border was deleted. *Hoxd* transcripts produced in E12.5 proximal limbs by the *Hox^DDel(attp-SB3)^* allele (Fig. 7A left, *Del(attp-SB3)/∆*) were scored by using both *in situ* hybridization and RT-qPCR (Fig. 7A right, S7A Fig. left). In such mutant proximal forelimb buds, *Hoxd11* to *Hoxd8* transcripts were depleted for more than 90% when compared to control proximal forelimbs (Fig. 7A right, S7A Fig. left). Surprisingly however, *Hoxd11* transcripts were not as dramatically affected in proximal mutant hindlimbs, and transcripts produced by the *Hoxd10* to *Hoxd*8 genes were decreased in amount by 50 to 60% when compared to control animals (Fig. 7A right, S7A Fig. left). The reduced level of *Hoxd* gene expression elicited by the mouse T-DOM deletion in the forelimb bud thus mimics the situation observed in chick proximal leg bud (S7A Fig. right) and these significant differences also exist in the way T-DOM operates in murine forelimb *versus* hindlimb buds.

**Fig. 7.**
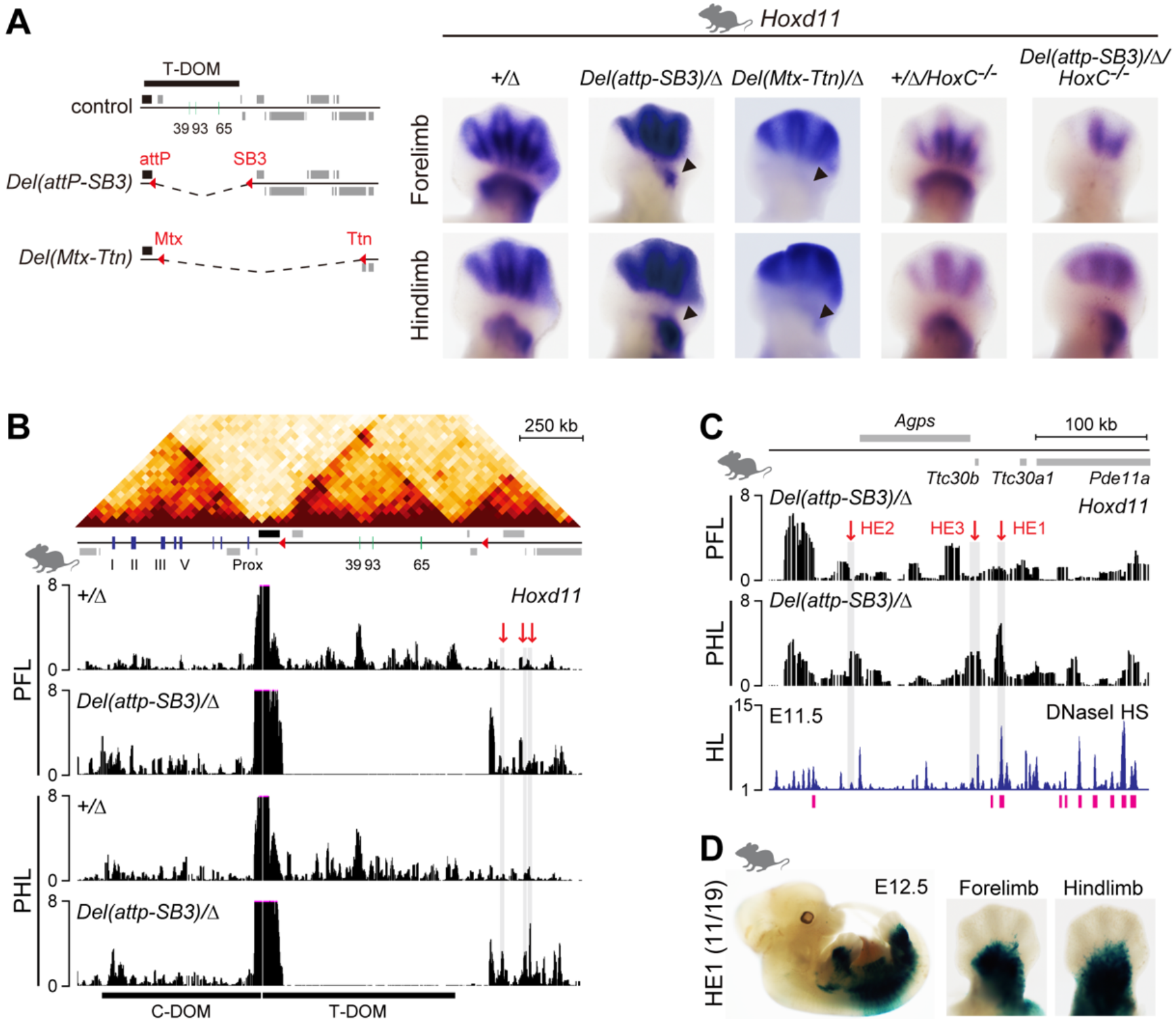
Different effects of a T-DOM deletion on fore- and hindlimb buds. (A) The *HoxD^Del(attp-SB3)^* and *HoxD^Del(Mtx-Ttn)^* alleles were constructed by deleting T-DOM through a ca. 1Mb or 2.1Mb large deletion, respectively (left, dashed line). *Hoxd11* expression in E12.5 fore- and hindlimbs from either control (*HoxD^Del(8-13)/+^*^)^ animals (indicated as ‘+/∆’) or mutants (*HoxD^Del(attp-SB3)/Del(8-13)^, HoxD^Del(Mtx-Ttn)/Del(8-13)^, HoxD^Del(8-13)/+;HoxC-/-^, HoxD^Del(attp-SB3)/Del(8-13)^;HoxC-/-*) littermates (indicated as ‘*Del(attp-SB3)/∆’*, ‘*Del(Mtx-Ttn)/∆’*, ‘*+/∆/HoxC-/-’*, ‘*Del(attp-SB3)/∆/HoxC-/-’*). In *Del(attp-SB3)/∆* mutant, *Hoxd11* expression was dramatically reduced in proximal forelimbs (arrowhead). Unexpectedly however, expression remained robust in proximal hindlimbs (arrowhead). In *HoxD^Del(Mtx-TiE2)^* mutant, *Hoxd11* expression disappeared in both proximal fore- and hindlimb buds (arrowhead). The absence of both T-DOM and the *HoxC* cluster did not affect *Hoxd11* expression. (B) HiC data adapted from ref[19] showing the two TADs on either side of the *HoxD* cluster and the TAD next to T-DOM. 4C profiles represent contacts established by *Hoxd11* in proximal fore- and hindlimb buds, from control or *Del(attp-SB3)/∆* mutant animals. In mutant cells deleted for T-DOM, (tracks 2 and 4), ectopic contacts between *Hoxd11* and regions in the neighboring TAD appeared due to their new proximity to *HoxD*. The shaded region with red arrows shows the domains where more contacts were detected in mutant proximal hindlimb versus proximal forelimb buds. (C) Enlargement of 4C profiles dispatched in (B), DNaseI, hyper-sensitive profiles using E11.5 embryos and potential limb enhancer regions (pink rectangles), as identified by using Limb-Enhancer Genie tool. Potential hindlimb-specific enhancers (HE) are shown by red arrows. (D) Mouse HE1 is active in the proximal fore- and hindlimb buds and in the trunk at E12.5. The number indicate stained embryos over total number of integrations. PFL, proximal forelimb; PHL, proximal hindlimb; HL, hindlimb.

Interestingly, the sustained expression of *Hoxd* genes in T-DOM deletion mutant proximal hindlimb buds completely disappeared when a larger deletion was engineered between the *Mtx* and *Titin (Ttn)* genes (Fig. 7A), indicating that the genomic regions between SB3 and *Ttn*, i.e. telomeric to the T-DOM TAD, contributes to the difference in gene expression observed between the mouse fore- and hindlimb buds when T-DOM is removed.

To identify potential forelimb to hindlimb differences in chromatin re-organization after the deletion of T-DOM, we generated 4C profiles from the mutant allele using the *Hoxd11* promoter as a viewpoint (Fig. 7B). In control proximal fore- and hindlimb cells, *Hoxd11* mostly contacted the intact T-DOM, with a particularly strong interaction with and around the CS39 region (Fig. 7B, tracks 1 and 3). In proximal cells deleted for T-DOM, interactions within the *HoxD* cluster were increased and ectopic contacts were established (or strongly re-enforced) with the newly fused neighboring telomeric TAD (Fig. 7B, tracks 2 and 4). We used this 4C-seq dataset to determine three candidate regions of potential enhancer activity, referred to as hindlimb enhancer (HE) 1 to 3 (Fig. 7B, C, red arrows). We cross-checked this selection either with DNaseI hypersensitive sites (HS) data from E11.5 hindlimb buds (GSM1014179)[53], with potential enhancer regions as defined by the Limb-Enhancer Genie tool[54] and with H3K4me1 ChIP-seq datasets obtained from control and mutant hindlimb proximal domains (Fig. 7C and S7B Fig). According to these criteria, HE1 turned out to be the most promising region and we thus assessed its enhancer potential in developing transgenic limb buds.

By using a transgenic enhancer reporter system, the HE1 region reproducibly drove *lacZ* expression in proximal fore- and hindlimb buds, lateral plate and somatic mesoderm at E12.5 (Fig. 7D). This result indicated that the HE1 enhancer activity is not specific for the proximal hindlimb, even though it was potentially active in a hindlimb-specific manner after deletion of T-DOM. To confirm this observation, we performed a 4C-seq analysis by using the HE1 sequence as a bait (S7C and S7D Fig), using both control and T-DOM deleted mutant proximal hindlimb cells. The 4C-profiles revealed that the HE1 sequence established slightly more contacts with the *Hoxd9* to *Hoxd12* DNA interval in mutant than in control hindlimb cells.

Finally, we looked at potential genetic interactions between the limb-specific differences in *Hoxd* gene expression and the *HoxC* gene cluster. Both *Hoxc10* and *Hoxc11* are indeed strongly transcribed in proximal cells of hindlimb buds (S7E Fig), whereas these transcripts were not detected in the equivalent forelimbs territories[55]. Furthermore, in proximal hindlimbs where *Hoxc* genes are expressed, the amount of *Hoxd* transcripts is decreased when compared to forelimb buds (Fig. 1C). Also, the deletion of *Hoxc11* on top of *Hoxa11/Hoxd11* double knockout mice exacerbated the observed hindlimb malformations[11][56], suggesting that HOXC proteins in hindlimb buds may help maintaining *Hoxd* transcripts, a function that would become visible in the absence of T-DOM, when *Hoxd* genes are no longer transcribed in forelimb proximal cells. We thus performed *in situ* hybridization for *Hoxd11* by deleting the entire *HoxC* cluster[57] on the top of the deletion of T-DOM. In these combined mutant limb buds, expression of *Hoxd11* was still detected in hindlimb proximal cells, indicating that the persistence of *Hoxd11* expression in hindlimb buds in the absence of T-DOM did not depend upon the presence of hindlimb proximal cells-specific *Hoxc* transcripts.

## DISCUSSION

### Conservation of the bimodal regulation in birds

While the expression of *Hox* genes belonging to the *HoxA*, *HoxC* and *HoxD* clusters during limb development are globally comparable between mammals and birds, some clear differences are nevertheless apparent, such as the quasi absence of *Hoxd* genes transcription in the proximal part of the developing leg buds in birds, while their expression there is required for proper mouse hindlimb development[56,58]. Also, while *Hoxd12* is expressed in the mouse limb buds like *Hoxd13*, i.e. mostly under the control of C-DOM only, its expression in the avian wing buds resembles that of *Hoxd11*, i.e. involving also the control of T-DOM, in future zeugopod cells. Because these expression specificities depend on the implementation of global regulations located within the two TADs flanking the *HoxD* cluster, we wondered whether the structures of these TADs were somehow modified in birds, or at least whether they would show some variations either between the two species, or between the bird wing and leg buds.

A global analysis of 4C and capture Hi-C datasets did not reveal any salient differences between mammals and birds regarding the way they implement this complex bimodal limb regulation at the *HoxD* locus. The TADs appeared well conserved between the two species, as well as the presence in chick of most- if not all-regulatory elements that had been described in the mouse counterparts, on both sides on the gene cluster[17,18], even though the chick TADs were reduced in size. We thus conclude that the bimodal regulatory strategy described in mammals (see[59]) is implemented in a similar manner during bird development. This re-enforces the idea that the function of *Hox* genes at these early steps of limb development is mostly to set up and organize the basic plan of the future appendages, rather than elaborating or fine-tuning on a pre-patterned structure.

### Interspecies comparison of the TAD boundary at *HoxD*

The bimodal regulatory mechanism and the transition between the two global controls at *HoxD* are thus well conserved amongst tetrapods. However, the distinct expression of *Hoxd12* in proximal limbs between mouse and chick suggests that the width of the TAD boundary at the *HoxD* locus may slightly vary between the two species. By using transcriptome, 4C and Hi-C datasets, we previously observed slightly different positions of this boundary in mouse distal *versus* proximal limb cells, due to the fact that *Hoxd10* and *Hoxd11* respond to both T-DOM and C-DOM regulations. We thus proposed that the boundary was located between *Hoxd11* and *Hoxd12* in proximal cells, while between *Hoxd9* and *Hoxd10* in distal cells[17,19] (Fig. 8A).

**Fig. 8.**
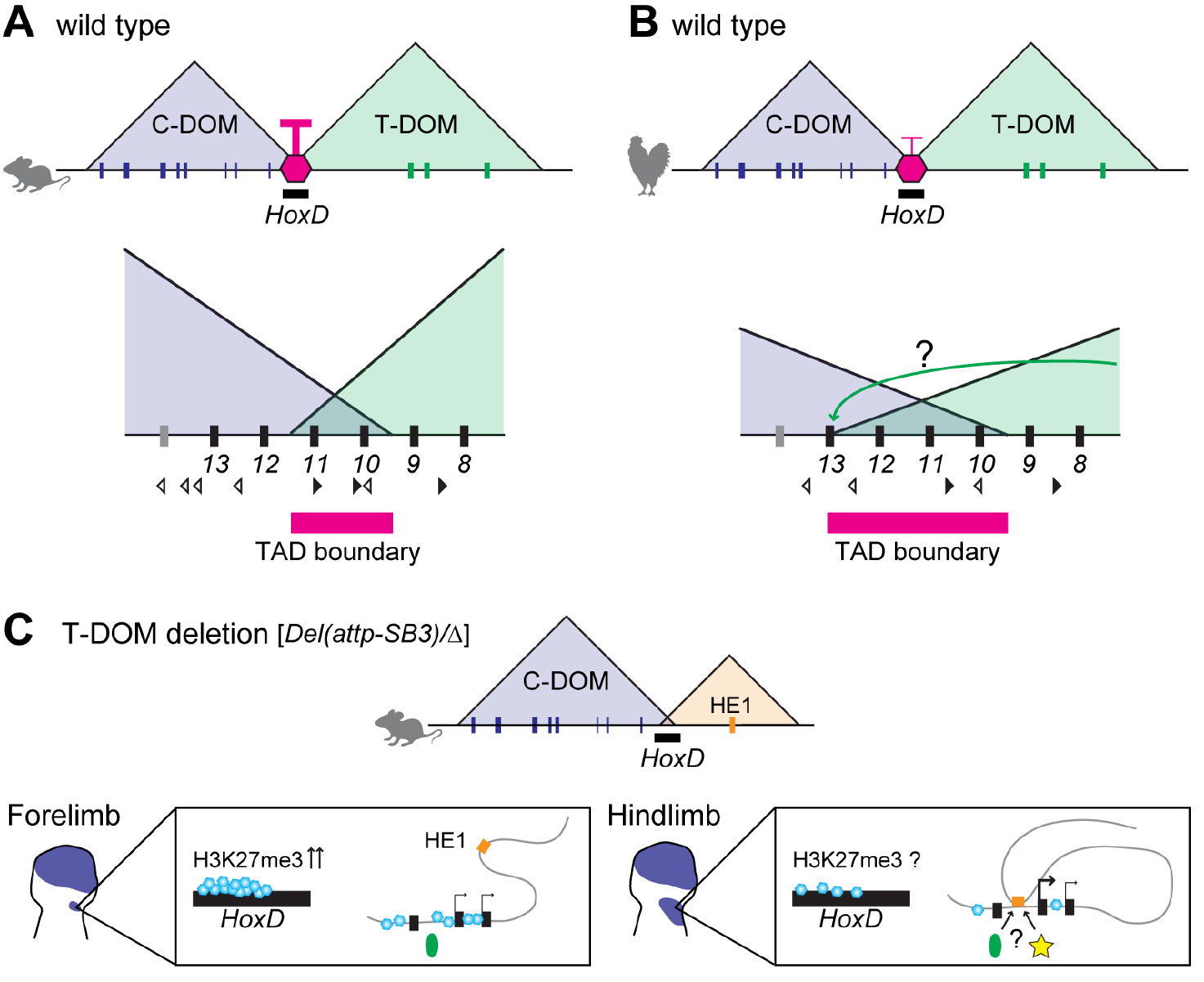
TAD boundaries at the mouse and chicken *HoxD* cluster and effect of T-DOM deletion. (A, B) TADs boundaries at the *HoxD* locus in mouse (A) and chick (B) limb buds. (A) In the mouse, the boundary moves along a few genes, in a window likely determined by a series of CTCF sites. In chick, the boundary appears less robust and slightly expanded towards the *Hoxd13* locus. This latter situation may enable T-DOM enhancers to interact with *Hoxd13* more efficiently in chick than in murine cells (B). Black and white arrowheads indicate the orientation of CTCF motives. (C) The different effects of T-DOM deletion in mouse proximal fore- and hindlimb cells. The transcriptional status of *Hoxd* genes is different between forelimb and hindlimb buds due to their prior expression or non-expression in lateral plate mesoderm, respectively. In the absence of T-DOM, H3K27me3 marks may accumulate over the silenced *Hoxd* genes in mutant forelimb buds as described in [17], making them poorly responsive to the HE1 enhancer. In contrast, mutant hindlimb bud proximal cells have their *Hoxd* genes poised for activation and *de novo* interactions with the HE1 enhancer may occur. H3K27me3 marks are in blue and presumptive factors, such as a potential mediator of DNA looping and a hindlimb-specific transcription factor, are shown by a green oval and a yellow star.

In contrast, the chick *Hoxd12* is strongly expressed in proximal wing buds, suggesting that the TAD boundary, in this context, expands towards the 5’ part of the gene cluster, close to the *Hoxd13* locus (Fig. 8B). Our CHi-C analysis re-enforced this view and positioned this boundary around the *Hoxd13* gene in chick limb buds at early stages (HH20), i.e. when T-DOM is active and controls the first phase of *Hoxd* genes transcription (S5 Fig). Subsequently (HH30), the boundary region was slightly reduced from *Hoxd13* to an area between *Hoxd13* and *Hoxd12* in chick limbs. Of note, *Hoxd12* is expressed in proximal limbs in geckos as in chicken[23], suggesting that the TAD boundary at the *HoxD* locus in proximal buds may have been slightly shifted during tetrapod evolution between birds and squamates on the one hand, and mammals on the other hand.

Different tetrapod species may thus display various positions of their TAD boundaries at *Hox* loci, acting as morphological cursors that would redistribute the various sub-sets of *Hox* genes responding to either proximal or distal enhancers. The distinct topologies of these boundaries may rely upon differences in the distribution and/or usage of CTCF binding sites. In the mouse, subsets of genes responding to either proximal or distal limb enhancers are delimited by different sets of bound CTCF sites [19] (Fig. 8A). Here, we show that chicken wing bud cells have less bound CTCF sites and of reduced intensity in the *HoxD* cluster than their murine counterparts, which could modulate the positioning of the boundary. In this case, the decrease in the overall strength of the boundary effect as a result of having less sites occupied by CTCF, may explain the extension of interactions up to *Hoxd12-Hoxd13* established by proximal enhancers (Fig. 8B). This hypothesis could nevertheless not be verified on chicken leg proximal cells as these cells do not strongly express the genes controlled by T-DOM.

### Distinct T-DOM regulations in mouse, chick and bat fore- and hindlimb buds

During bat limb development, *Hoxd10* and *Hoxd11* transcripts are progressively lost throughout the hindlimbs only, in part due to the distinct enhancer activity of the BAR116 sequence located within T-DOM[6]. When the mouse BAR116 cognate sequence was used in a transgenic context, no activity was detected in any limb cells. Likewise, when we used mouse *CS9*, i.e. a shorter fragment of the *CS93* sequence, staining was not observed. However, when the full length *CS93* sequence was injected, a robust enhancer activity was scored in a proximal limb region (Fig. 3). This discrepancy between two experiments involving almost the same sequences may be caused by the positions of the regions used for the mouse transgenic enhancer assays, the mouse sequence extending a bit more towards one of its extremities. Either the enhancer activity was provided by this sub-fragment, or this fragment may be required for the expression of a more widespread activity of the full DNA sequence. It remains that the BAR116 sequence may not be a bat specific enhancer.

However, while the bat BAR116 showed different enhancer activities between the mouse transgenic fore- and hindlimb buds, i.e. strong in forelimb and weak in hindlimb, the mouse equivalent appeared to have similar enhancer activities between fore- and hindlimbs, in agreement with the continuous expressions of *Hoxd10* and *Hoxd11* in both types of limbs. To further validate this correspondence, we looked at the behavior of the chick *CS93* sequence in the same mouse transgenic context. While two sets of patterns were obtained with various distal to proximal distributions of the *lacZ* staining, a clear imbalance was scored between forelimb and hindlimb cells, with a much stronger expression in the former than in the latter. Therefore, the chick enhancer sequence behaved more like the bat sequence than like their murine counterparts, an effect that was re-enforced by the sequence alignments, which revealed more similarities between chick and bats than between the two mammalian species. This similarity correlates with *Hoxd* gene expression and may relate to the salient distinctions in morphologies between fore- and hindlimbs.

### Premature termination of T-DOM regulation in chick hindlimb buds

The termination of the T-DOM regulatory activity in proximal limb cells coincides with the binding of the HOXA13 protein at various sites within the TAD. Also, the removal of both *Hoxa13* and *Hoxd13* functions lead to the continuation of T-DOM regulation and to the failure of C-DOM activation, suggesting that HOX13 proteins are necessary to terminate T-DOM function and to implement the bimodal switch[20,49]. The chick *Hoxd13* gene starts to be expressed at around stage HH18-19[31] when H3K27ac enrichment is not yet detected over C-DOM (except for *island I*)(Fig. 4). Instead, H3K27me3 marks are still present over C-DOM at this early stage, unlike in the early mouse limb buds (Fig. 4 and S3 Fig)[17], suggesting that *Hoxd13* early activation in chick may be driven by the T-DOM regulation until the C-DOM regulation is implemented and takes it over. This idea is well supported by our CHi-C analysis showing that the TAD boundary is expanded towards the *Hoxd13* locus in the early chick limb buds.

In addition, a major difference was observed in the activation of *Hoxa13* between chick and mouse hindlimb buds, with an earlier and stronger activation of *Hoxa13* in chick leg buds at HH20, when compared to both mouse hindlimb buds and chick wing buds. This indicates that the T-DOM activity may be readily terminated by the early presence of HOXA13 protein and that the C-DOM regulation is ready to be active earlier in chick leg buds than in wing buds. The potential causes for both this early activation of *Hoxa13* in chick leg buds and the strong level of H3K27me3 observed over C-DOM in chick wing and leg buds remain to be determined.

### Enhancer reallocation and anterior-posterior (AP) position of the limb buds

In the mouse, we show that the deletion of the telomeric TAD has different effects upon forelimb and hindlimb proximal cells, as fair levels of *Hoxd* gene expression remain only in the proximal hindlimb domain. Since a larger deletion including more telomeric sequences totally abrogated *Hoxd* expression in proximal hindlimb, we concluded that additional hindlimb-specific enhancers may be located telomeric of the T-DOM TAD. Interaction profiles established after the deletion of T-DOM revealed novel hindlimb-specific contacts between target *Hoxd* genes and the newly identified HE1 sequence, which is normally located outside T-DOM but brought to the vicinity of the cluster after the deletion. The HE1 region is located near the *Agps* and *Pde11a* genes and our 4C profile using HE1 as a bait showed strong interactions with both genes under normal condition. Noteworthy, *Agps* is involved in the rhizomelic chondrodysplasia punctate 3 (RCDP3) condition, with a shortening of proximal limbs[60,61], suggesting that HE1 may be involved in the regulation of the *Agps* gene. The deletion of T-DOM may thus reallocate part of the HE1 proximal limb enhancer activity towards *Hoxd* promoters.

Our genetic approach however makes it difficult to assess whether this sequence is used for *Hoxd* regulation under normal circumstances, or whether the interactions observed are mostly due to its new proximity to the target genes induced by the deletion of T-DOM. In the former case, this may indicate that as in chick and bats, the global C-DOM regulation may be more active in forelimb than in hindlimb buds and hence the HE1 enhancer may not be necessary. In the case of the mouse, this deficit of regulation during proximal hindlimb development could have been compensated for by evolving additional enhancers outside the TAD (Fig. 8C). The HE1 sequence is bound by several factors, such as the YY1, proposed to mediate enhancer-promoter contacts at distance in ESCs[62], or PITX1, a hindlimb-specific factor[63], suggesting that multiple transcriptional factors may contribute to this interaction.

Finally, the sustained expression of *Hoxd* genes observed in T-DOM deleted mutant proximal hindlimb cells may merely reflect the history of early limb bud cells. In the wild type condition indeed, the anterior bud emerges from a field of lateral plate mesoderm (LPM) devoid of transcripts for *Hoxd9*, *Hoxd10* or *Hoxd11*. In contrast, posterior limb buds derive from LPM cells already expressing these genes, due to their more posterior AP position along the trunk mesoderm. In the absence of T-DOM, expression of these genes would not occur in the anterior buds due to their repressed state and the lack of appropriate enhancers, whereas expression could be inherited and maintained in the posterior buds through a mechanism independent of T-DOM, perhaps involving the HE1 sequence.

## Materials and Methods

### Animal experimentation

All experiments involving animals were performed in agreement with the Swiss law on animal protection (LPA), under license No GE 81/14 (to DD). Chick embryos from a white Leghorn strain were incubated at 37.5°C and staged according to ref[51].

### *In situ* hybridization and *lacZ* staining

Whole mount *in situ* hybridization was performed as described previously[64]. For *lacZ* staining, embryos were fixed in 1x PBS pH7.39-7.41, 2mM MgCl_2_, 4% PFA/PBS, 0.2% glutaraldehyde and 5mM EDTA for 20 min at room temperature and washed three times for 20 min in 1x PBS, 2mM MgCl_2_, 0.2% NP40 and 0.01% sodium deoxycholate. Samples were stained in 5mM potassium ferrocyanide, 5mM potassium ferricyanide and 0.5 mg/ml X-gal at room temperature overnight, followed by wash with the wash solution once and refixation with 4%PFA/PBS at 4°C overnight.

### Micro-dissection chick limbs at HH30

Limb tissues at HH30 were micro-dissected into acropod (distal), mesopod and zeugopod (proximal) regions, respectively. Only distal and proximal regions were used for RNA-seq and 4C-seq.

### RNA-seq and Data analysis

Total RNA was extracted from mouse and chick limb bud tissues using the RNeasy Micro Kit (QIAGEN) following the manufacturer instructions. Libraries were prepared with at least 200 ng of total RNA following Illumina TruSeq stranded mRNA sample preparation guide. Sequencings were performed with 100-bp or 75-bp single-end. The data was mapped onto either GRCm38 (mm10) or the International Chicken Genome Reference Consortium Gallus_gallus-5.0 (galGal5) using the Tophat2 (Version 2.0.9)[65] and unique mapped reads were extracted. The number of uniquely mapped reads was calculated using FLAGSTAT (SAMtools, Version 0.1.18)[66] and this value was used for the subsequent normalization of each coverage data to be the million reads number. All analyses were processed by our Galaxy server (the Bioteam Appliance Galaxy Edition, https://bioteam.net, https://bioteam.net/products/galaxy-appliance)[67].

### RNA extraction and RT-qPCR

Total RNA was extracted using the RNeasy Micro Kit (QIAGEN), following the manufacturer instructions. 1 μg total RNA was used for cDNA synthesis with SuperScript VILO (Invitrogen). RT-qPCR was performed on a CFX96 real-time system (BIORAD) using the GoTaq qPCR Master Mix (Promega). Each RT-qPCR was carried out with at least two biological replicates and each experimental information is described in S1 Table. Primer sequences for qPCR are listed in S2 Table.

### 4C-seq and Data analysis

Each mouse and chick limb tissue were fixed separately with 2% formaldehyde, lysed and stored at −80°C. Samples were digested with *NlaIII* and *DpnII* as primary and secondary restriction enzymes, respectively, and ligation steps were performed using high concentrated T4 DNA ligase (Promega)[68]. Inverse PCRs for amplification were carried out using primers for each viewpoint[69](S3 Table). PCR products were multiplexed and sequenced with 100bp single-end, followed by post-processing (de-multiplexing, mapping, and 4C analysis) using the HTS station (http://htsstation.epfl.ch)[70]. Fragment scores were normalized to the mean score of fragments falling into a region defined as the bait coordinated ± 1 Mb, and the data were smoothed by using a running mean with a window size of 11 fragments. The information regarding fragments excluded during the procedure is provided in S3 Table. A relative quantification of the signal spanning both the *HoxD* telomeric or centromeric domains was performed. Genomic coordinates for calculations are:

mm10, chr2: 73,921,943-74,648,943 / chr2: 74,765,943-75,601,943;
galGal5, chr7: 15,920,642-16,318,067 / chr7: 16,414,183-16,699,172.

This quantification is not absolute and only reflects the balance of contacts between the two domains, for each sample.

### ChIP-seq and Data analysis

Chromatin immunoprecipitation experiments were performed as previously described[20]. Micro-dissected limb tissues from mouse and chick embryos were cross-linked with 1% formaldehyde/PBS for 15 min at room temperature. Chromatin was sheared and used for each immunoprecipitation with anti-H3K27ac (ab4729, Abcam), anti-H3K27me3 (07-449, Merck Millipore), anti-H3K4me1 (ab8895, Abcam) and anti-CTCF (61311, Active Motif). Libraries were prepared with at least 5 ng of purified DNA following Illumina TruSeq ChIP library preparation guide. Sequencing was performed with 100bp single-end. Demultiplexed ChIP-seq reads were mapped onto the galGal5 or mm10 using Bowtie (Version 0.12.7)[71], with parameters “-m1 –strata –best” according to conditions described previously[72], and PCR duplicates were removed from mapped reads using SAMtools (Version 0.1.18)[66]. By using bamCompare (Version 2.5.0), the ChIP data from H3K27ac, H3K27me3 and H3K4me1 and the input data were normalized and compared to compute the log2 of the number of reads ratio. The CTCF ChIP data was normalized to obtain 1x depth of coverage by using bamCoverage (Version 2.5.0)[73,74]. The CTCF motif orientation analysis was performed as previously described[19]. All analysis was done with our Galaxy server (the Bioteam Appliance Galaxy Edition, https://bioteam.net, https://bioteam.net/products/galaxy-appliance)[67].

### SureSelect probe design and Capture Hi-C (CHi-C)

The library of SureSelect enrichment probes were designed over the genomic interval (galGal3: chr7:15,990,001-19,170,000) using the SureDesign online tool of Agilent. Probes cover the entire genomic region (galGal5, chr.7: 14,946,000-17,870,000) and were not designed specifically in proximity of *DpnII* sites. Dissected tissues were dissociated in 10% FCS/PBS with collagenase (C7657, Sigma) to a final concentration of about 1.3 μg/μL, and samples were incubated in Thermomixer at 37°C at 800 rpm for 20 min. After discarding the supernatant, cells were crosslinked with 1% formaldehyde/PBS at RT for 10 min, quenched with glycine and centrifuged to discard the supernatant. Cells were resuspended with PBS containing proteinase inhibitor and then centrifuged again. After removing supernatant, cells were kept at −80°C before use. Hi-C library preparation was performed as described in[75], with the following changes: (i) Resuspended crosslinked cells in ice-cold Lysis buffer were placed on a rotation wheel at 4°C at 30 rpm for 30 min for cell lysis. (ii) For chromatin digestion, 400U of *DpnII* (R0543M, New England Biolabs) were added to the samples and incubated at 37°C at 700 rpm for 4 hrs. Another 400U of *DpnII* were added and samples were incubated overnight. (iii) Blunt-end ligation of biotin filled-in DNA was performed at RT at 30 rpm on rotating wheel for 4 hrs. (iv) No removal of biotin from un-ligated ends was performed. (v) DNA was sheared to a size of 200 to 800bp by using COVARIS E220, with the following conditions; 175W, 10% Duty factor, 200 Cycles per Burst, 60 sec. (vi) DNA pulldown was performed by using Dynabeads MyOne Streptavidin T1 (65601, Thermo Fisher). (vii) DNA was measured by Qubit and 200ng were used for further treatment followed by manufacturer’s protocol (SureSelect^XT^ Target Enrichment System for Illumina Paired-End Multiplexed Sequencing Library).

### CHi-C data analysis

Paired-end sequencing data were processed as follows. First, adapters were removed using cutadapt version 1.6[76] with the following parameters: -a AGATCGGAAGAGCACACGTCTGAACTCCAGTCAC for R1 and -a AGATCGGAAGAGCGTCGTGTAGGGAAAGAGTGTAGATCTCGGTGGTCGCCGTATCATT for R2. They were then processed by using hicup version 6.1.0 with the bowtie2 version 2.2.6[77] and SAMtools version 1.2[66], with galGal5 as reference genome and GATC as restriction enzyme recognition sequence. The pairs were next converted from bam to tabulated files, with the position of the middle of the fragment to which hicup assigned the read, by using an ad-hoc python script (available upon request).

Only valid pairs with both MAPQ above 30 were kept. Then, pairs with both mates in the capture region (galGal5, chr7:14,946,000-17,870,000) were extracted and processed with cooler to obtain a balance matrix of the capture region with 5kb bins. The Fig. 5 and S5 Fig were obtained with personal R scripts (available upon request) (http://www.R-project.org). S5A and S5B Fig are the balanced matrices with linear scale. Fig 5 was obtained by subtracting the two balanced matrices. To assess the significance of increased contact between two regions, a Wilcoxon rank sum test was performed using R with the values of the bins in the region of the two balanced matrices. Because 75% of valid pairs MAPQ30 do not involve the capture region, all valid pairs were also processed with cooler to obtain a balance matrix of the whole chromosome 2 at 40kb. These matrices were used in S6 Fig. To define TAD borders (Fig 5), the TopDom algorithm[42] was run with a window size of 28 from the 10kb binned balanced matrices, as gaps were too numerous at a 5kb resolution.

### Mutant stocks

The *HoxD^Del(8-13)^* and *HoxD^Del(attp-SB3)^* alleles have been previously described[17,78]. The *HoxD^Del(Mtx-Ttn)^* allele was produced by TAMERE using the TiE2 (*Titin* exon 2) allele[79] (kindly provided by Dr. Michael Gotthardt) and a *Mtx2* gene trap allele (https://igtc.org/cgi-bin/annotation.py?cellline=CSI574). The sequences of genotyping primers are indicated in S2 Table. All embryos analyzed in Fig 7 and S7 Fig were heterozygotes, balanced by the *HoxD^Del(8-13)^* allele.

### Analysis of sequence alignment and limb enhancer prediction

mVista, tools for comparative genomics was used for comparison between sequences of the mouse *CS93* (mm10, chr2: 75,208,103-75,210,328), the bat BAR116 (Myoluc2, GL429772: 6,606,808-6,608,652) and the 2kb region containing the chick *CS93* (galGal5, chr7: 16,104,863-16,106,863), with LAGAN alignment program with default parameter (http://genome.lbl.gov/vista/index.shtml). Potential limb enhancer regions were identified by using Limb-Enhancer Genie tool, with the following condition: (i) analysis type: Scan for top, (ii) method: Combined Model (https://leg.lbl.gov/)[54].

### Enhancer transgenic assays

For the enhancer assays, embryos carrying the mouse *CS93/lacZ* and *HE1/lacZ* were generated by lentivirus-mediated transgenesis and pronuclear injection, respectively. The mouse *CS93* (mm10, chr2: 75,208,104-75,210,328) was amplified from C57BL/6 genomic DNA and cloned into the *pRRL-lacZ* vector, as described previously[17]. Lentiviruses were produced and injected into the peri-vitelline space of mouse zygotes[32]. The mouse *HE1* (mm10, chr2: 75,959,179-75,960,378) and the region containing the chick *CS93* sequence (galGal5, chr7: 16,104,863-16,106,863) were obtained from B6CBAF1/J and White Leghorn genomic DNA, respectively, and cloned into a *βglobin-lacZ* vector. The construct was injected into mouse oocytes. All transgenic embryos were harvested between E10.5 and E12.5 and used for *lacZ* staining.

### Accession Numbers

RNA-seq, 4C-seq, ChIP-seq and CHi-C datasets are available from the NCBI Gene Expression Omnibus repository under accession number GSE115563. The public datasets used in this research are listed in S4 Table.

## ACKNOWLEDGEMENTS

We thank Dr. A. Necsulea for help with bioinformatic analyses, S. Gitto and T.-H. Nguyen Huynh for technical help, Drs. J.-M. Matter, M. Gotthardt, M. Ros, C. Tabin, Y. Kawakami and K. Tamura for sharing materials, the Geneva Genomics Platform (University of Geneva), the transgenic core facilities (University of Geneva and Ecole Polytechnique Fédérale in Lausanne), the Gene Expression Core Facility, the Bioinformatics and Biostatistics Core Facility of the Ecole Polytechnique Fédérale in Lausanne and the ENCODE Consortium and the ENCODE production laboratory generating the particular dataset. We also thank all members of the Duboule laboratories for discussions.

## Author Contributions

Conceptualization: N.Y.-K., G.A. and D.D.

Methodology: N.Y.-K., G.A., C.C.B. and D.D.

Investigation: N.Y.-K., C.C.B., G.A. and L.B.

Formal Analysis: N.Y.-K. and L.L.-D.

Writing-Original Draft: N.Y.-K., L.L.-D. and D.D.

Writing-Review and Editing: L.L.-D., G.A., L.B. and C.C.B.

Supervision and funding: D.D.

## SUPPORTING INFORMATION

**S1 Fig (related to Fig 1).**
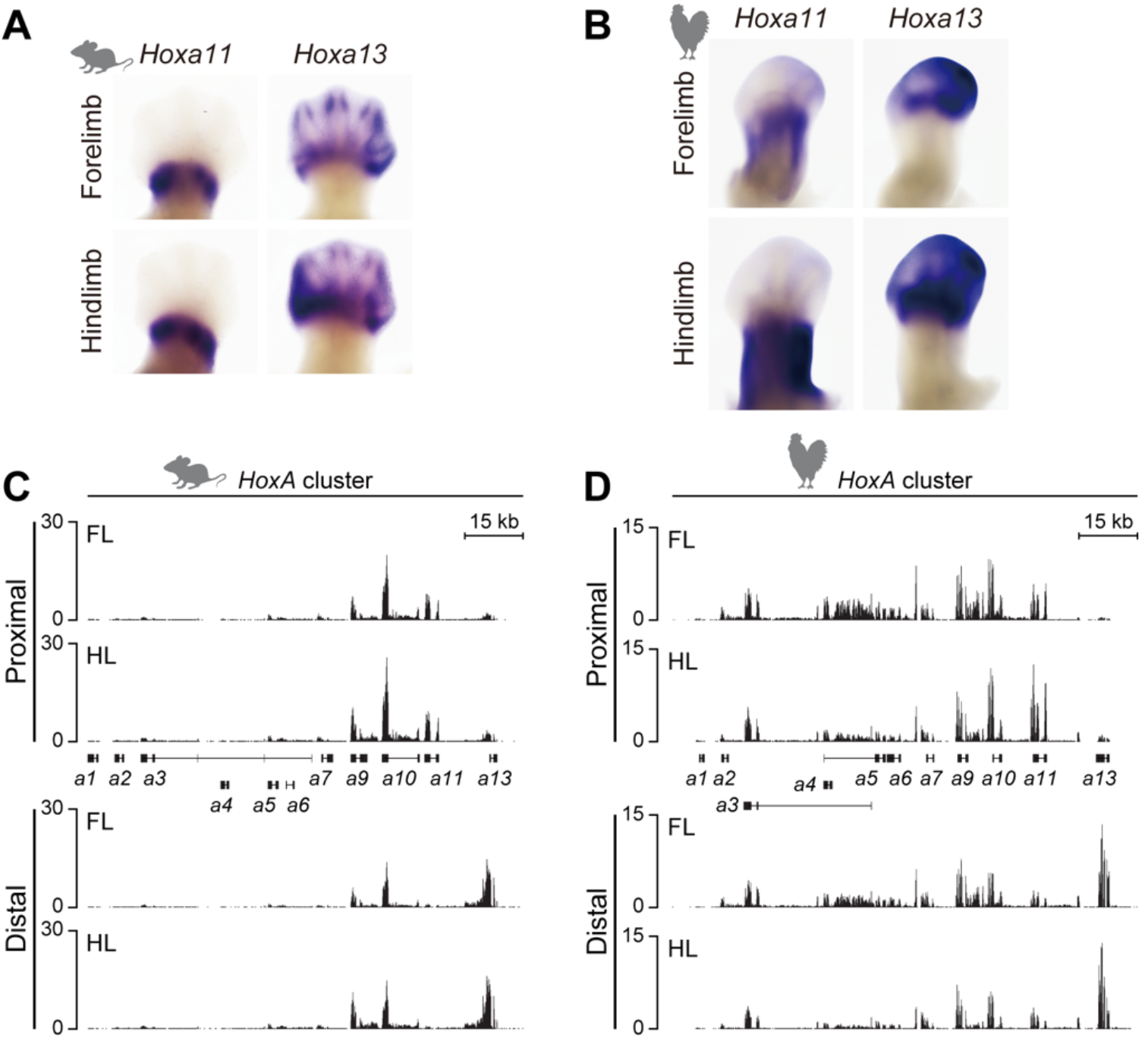
*Hoxa* genes expression in mouse and chick limb buds. (A, B) *In situ* hybridization analysis of E12.5 mouse and HH28 chick fore-and hindlimb buds showing the expression of *Hoxa* genes. (A) Expression patterns of *Hoxa11* and *Hoxa13* in mouse forelimb are similar to those in hindlimb at E12.5. (B) Stronger expression of *Hoxa11* is observed in the chick leg bud than in the wing bud. (C, D) Transcription profiles of *Hoxa* genes in micro-dissected proximal and distal domains from either E12.5 mouse (C) or HH30 chick (D) fore-and hindlimb buds. Right limbs in (A, are oriented with proximal to the bottom and distal to the top. FL, forelimb; HL, hindlimb. The *Y* axis represents the strand-specific RNA-seq read counts, normalized by the total number of million mapped reads.

**S2 Fig (related to Fig 2).**
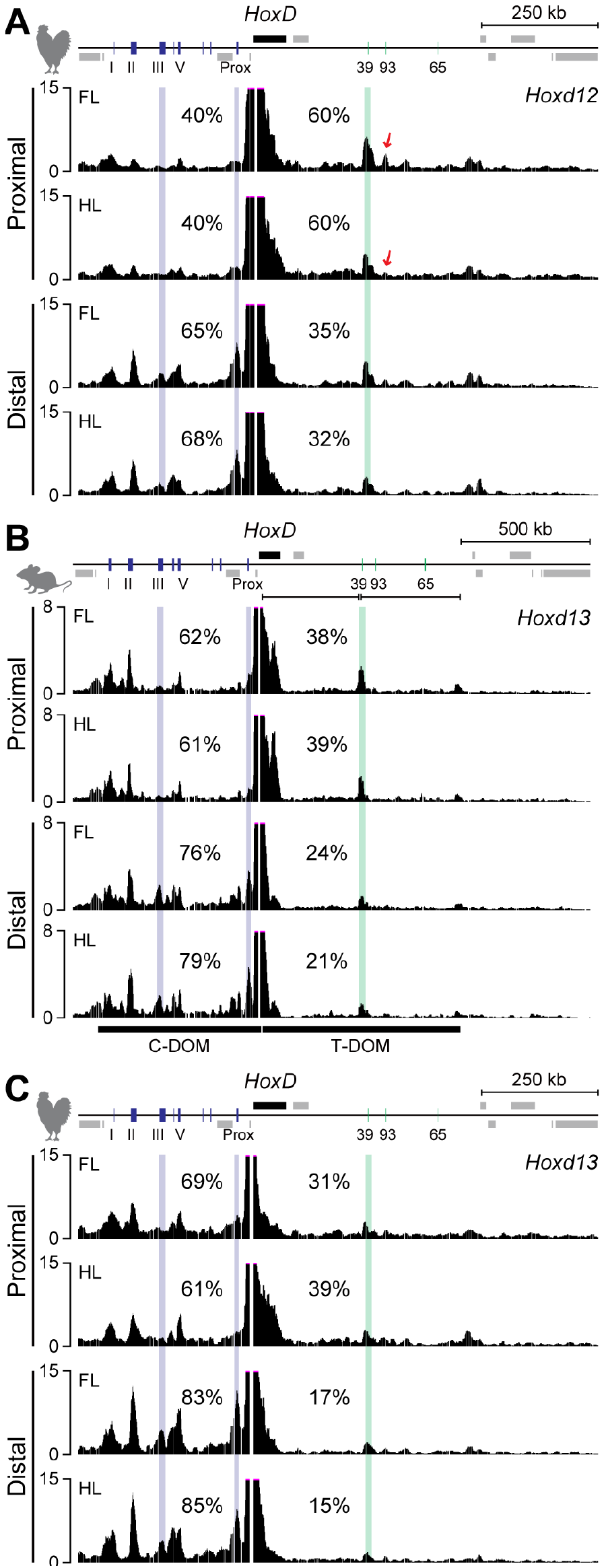
The regulatory switch between TADs in mouse and chick limb buds. (A-C) The 4C interaction profiles with chick *Hoxd12* (A), mouse *Hoxd13* (B) and chick *Hoxd13* (C), in mouse (E12.5) and chick (HH30) fore-and hindlimbs. (A) In addition to the *CS93* region, contacts between *Hoxd12* and the *CS39* region were also reduced in chick proximal leg bud cells. In the distal wing-and leg bud cells, *Hoxd12* mainly contacted C-DOM, in contrast to the profile observed with the *Hoxd10-11* bait. (B, C) Both mouse *Hoxd13* and chick *Hoxd13* promoters constitutively interacted with C-DOM. The interaction between *Hoxd13* and either island III or *Prox* specifically increased in both mouse and chick distal limbs. FL, forelimb; HL, hindlimb.

**S3 Fig (related to Fig 4).**
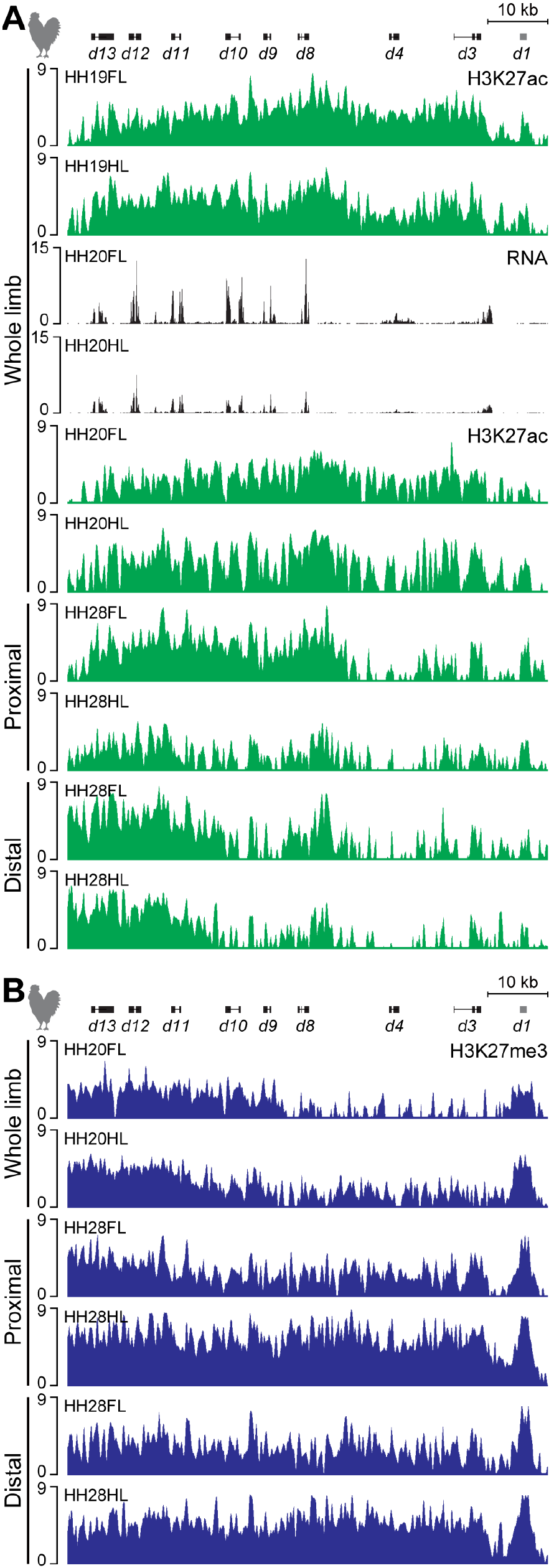
H3K27ac, H3K27me3 and RNA-seq at *HoxD* in chick limbs. (A) H3K27ac marks (tracks 1-2 and 5-10) and transcription profiles (tracks 3 and 4) at the *HoxD* locus either in whole, proximal or distal wing-and leg buds. 5’ *Hoxd* genes are covered by H3K27ac in the leg bud at HH19 and HH20. However, the level of *Hoxd* transcripts was reduced at HH20 (see also S3B Fig, track 4). In proximal leg buds at HH28, a significant decrease in H3K27ac enrichment was detected, which corresponded to the reduction in *Hoxd* expression (track 8). (B) H3K27me3 distribution in either whole, proximal or distal wing-and leg buds at HH20 and HH28. Stronger enrichments were observed in both whole leg buds at HH20 and proximal leg buds at HH28, when compared to the corresponding samples from wing buds. The *Y* axis represents the strand-specific RNA-seq read counts, normalized by the total number of million mapped reads. Enrichment (*Y-axis*) of ChIP is shown as the log_2_ ratio of the normalized number of reads between ChIP and input samples. FL, forelimb; HL, hindlimb.

**S4 Fig (related to Fig 4).**
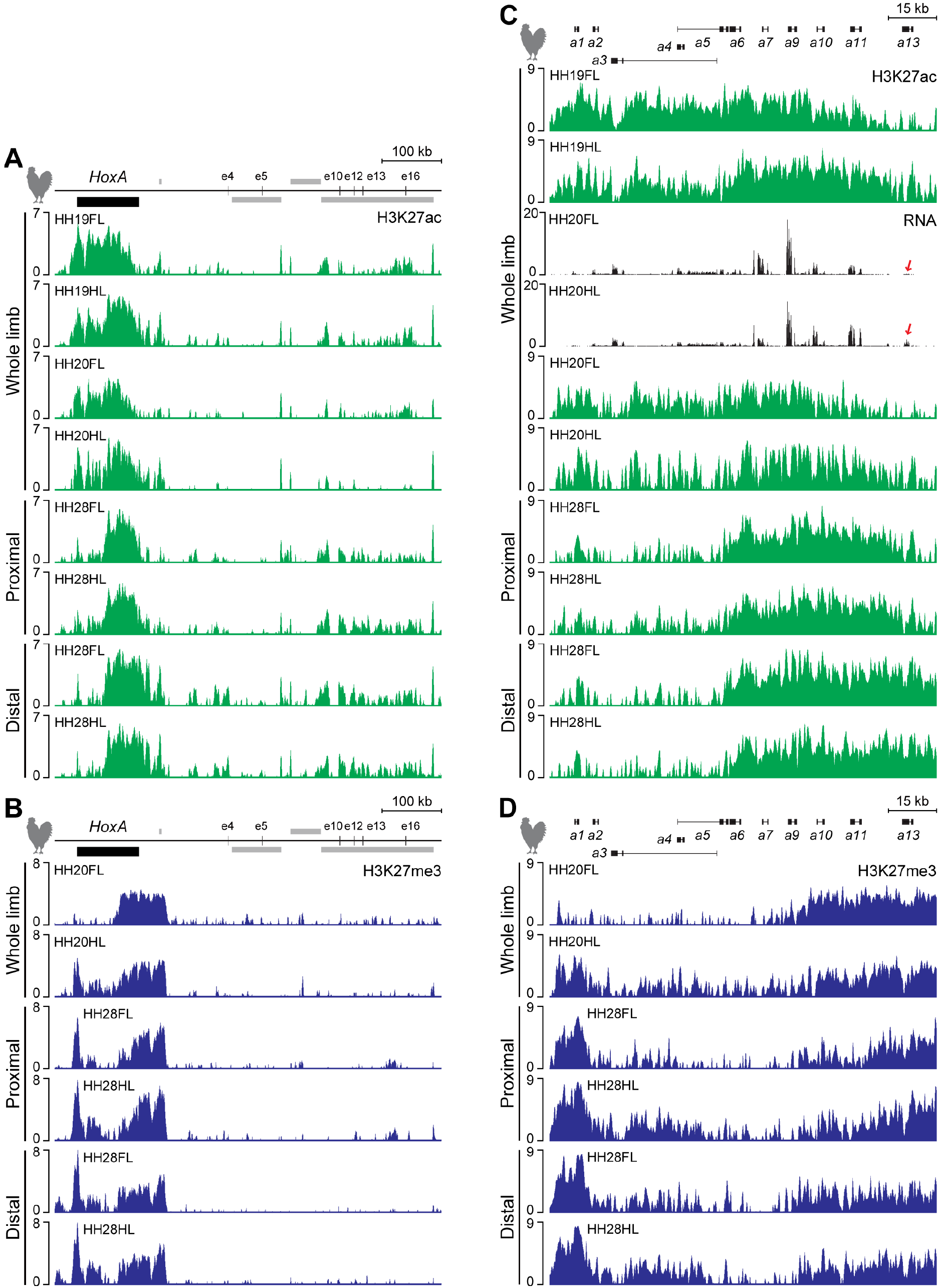
H3K27ac and H3K27me3 profiles of ChIP-seq and RNA-seq at the *HoxA* locus in chick limbs. (A, B) Distributions of H3K27ac and H3K27me3 marks over the *HoxA* cluster and its regulatory elements in either whole, proximal or distal wing-and leg buds at HH19, HH20 and HH28. (A) Stronger enrichment of H3K27ac around the 5’ *Hoxa* genes were observed in leg buds at both HH19 and HH20, whereas less marks were scored at HH20, in the region covering the *e10* to *e16* enhancers, when compared to wing-and leg buds at HH19. At HH28, profiles established from proximal or distal region were comparable between wing-and leg buds. (B) 3’ *Hoxa* promoters were not labelled by H3K27me3 marks in wing buds at HH20 (track 1). Strong enrichments of H3K27me3 over the *HoxA* regulatory elements were not scored, unlike in both C-DOM and T-DOM at the *HoxD* locus (see also Fig. 4B). (C) H3K27ac marks (tracks 1-2 and 5-10) and transcription profiles (tracks 3 and 4) at the *HoxA* locus in either whole, proximal or distal wing-and leg buds. More H3K27ac marks were detected at 5’ *Hoxa* genes in whole leg buds at both HH19 and HH20, corresponding to higher levels of *Hoxa* gene transcripts in leg buds than in wing buds (red arrows in tracks 3 and 4). (D) H3K27me3 profiles in either whole, proximal or distal wing-and leg buds at HH20 and HH28. The *HoxA* regulatory elements at the chick locus were identified by using mouse coordinates and the LiftOver function of the UCSC genome browser. The *Y* axis represents the strand-specific RNA-seq read counts, normalized by the total number of million mapped reads. Enrichment (*Y-axis*) of ChIP is shown as the log_2_ ratio of the normalized number of reads between ChIP and input samples. FL, forelimb; HL, hindlimb.

**S5 Fig (related to Fig 5).**
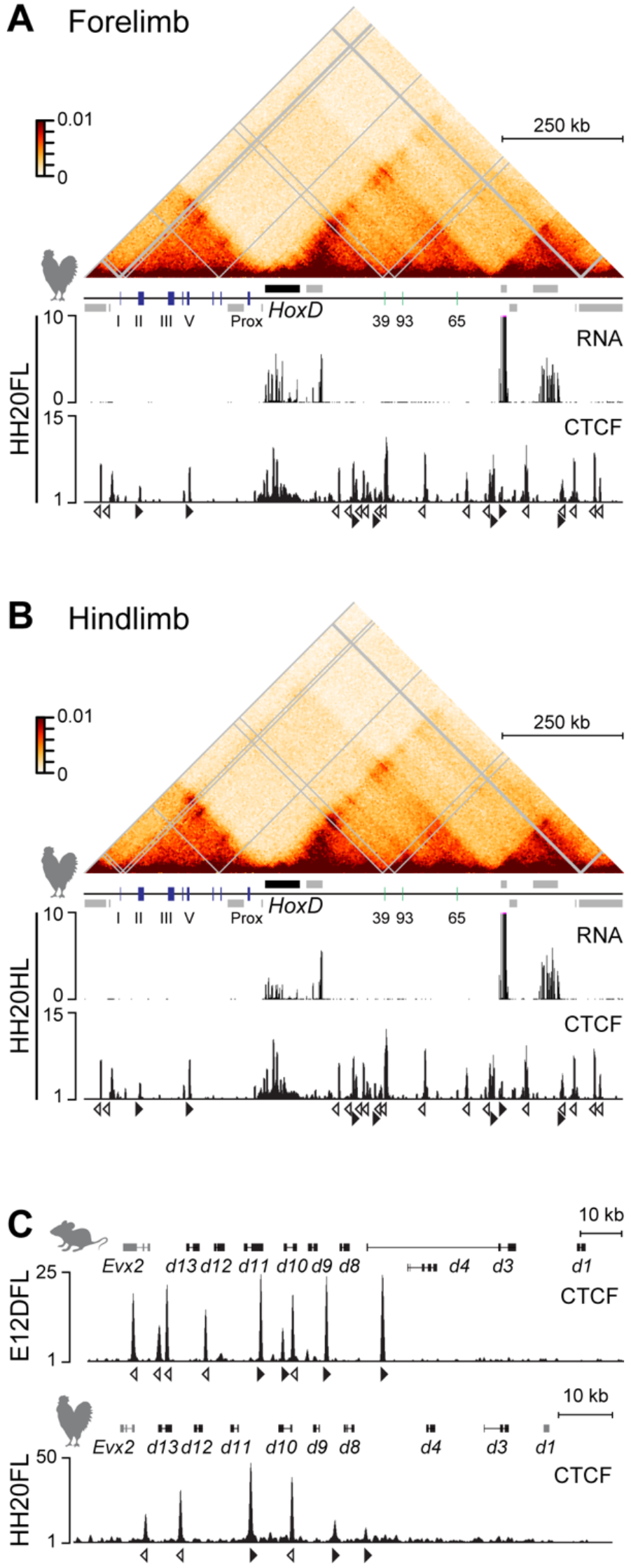
Chromatin conformation at the chick *HoxD* locus in wing-and leg buds and conservation of CTCF sites orientations. (A, B) Capture Hi-C (CHi-C) heat maps at 5kb resolution, transcription profiles and CTCF ChIP-seq by using either whole wing-or leg buds at HH20. Both CHi-C heat maps and CTCF distributions were relatively similar between wing-and leg buds, although some slight differences were detected. A noticeable downregulation of *Hoxd* gene expression was observed in leg buds when compared to wing buds. (C) Comparison of CTCF sites orientations at the *HoxD* cluster between mouse distal forelimb at E12.5 (top) and chick wing bud at HH20 (bottom). Opened and closed arrowheads indicate the orientation of the CTCF motives. The *Y* axis represents the strand-specific RNA-seq read counts, normalized by the total number of million mapped reads. Enrichment (*Y-*axis) is shown at the normalized 1x sequencing depth of CTCF ChIP. FL, forelimb; HL, hindlimb.

**S6 Fig (related to Fig 6).**
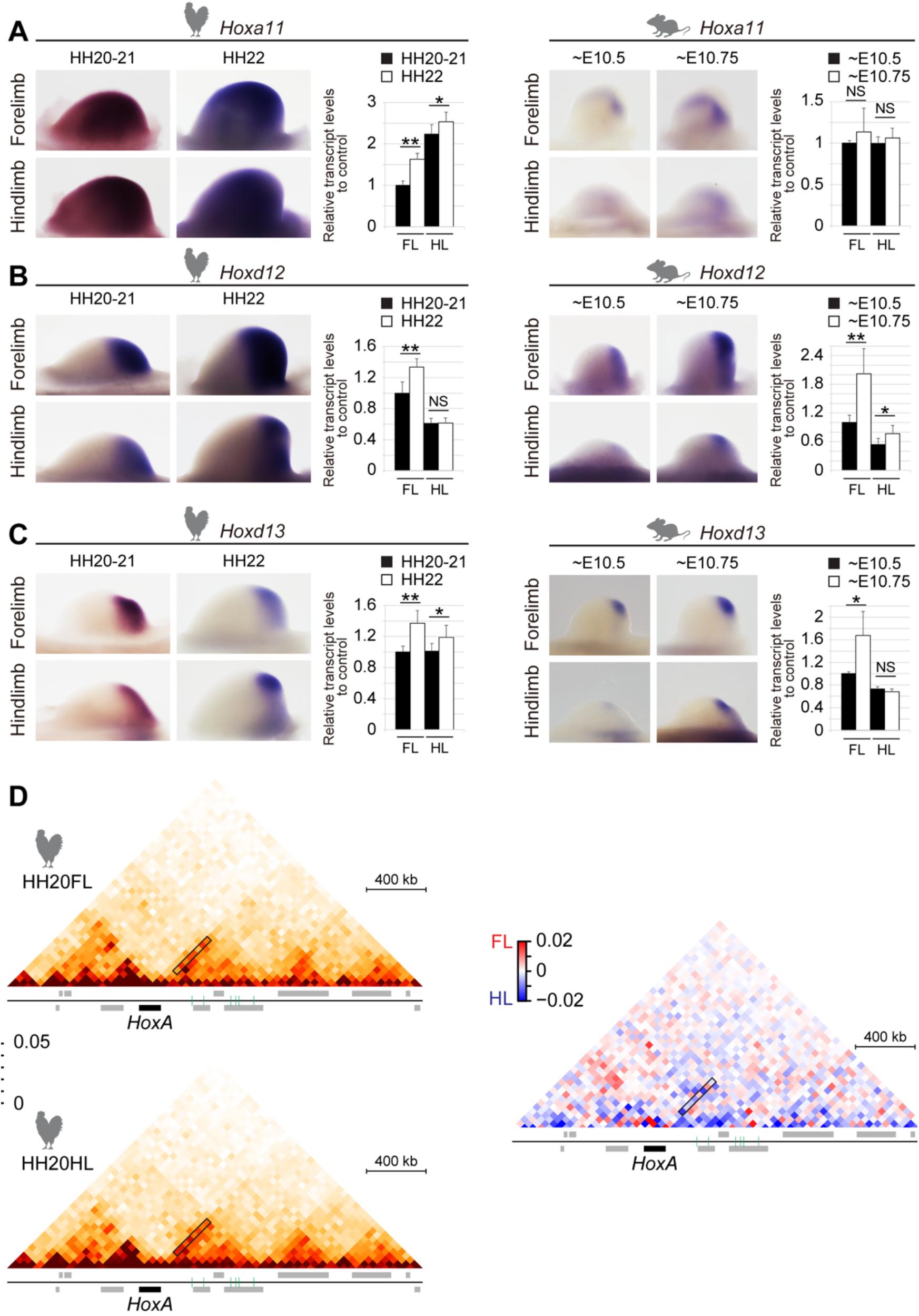
Expression of *Hoxa* and *Hoxd* genes in chick and mouse limb buds. (A) *Hoxa11* expression was stronger in chick leg buds than in wing buds (left), whereas it seems increased in both mouse fore-and hindlimb buds, as development proceeded (right). (B) Expression of *Hoxd12* in both chick wing buds and mouse limb buds displayed a similar trend, except for the chick leg buds. (C) Expression of *Hoxd13* in both chick limb buds and mouse forelimb buds were similar, except in mouse hindlimb buds. (D) Hi-C data at *HoxA* with 40kb resolution using wing-and leg buds at HH20. More contacts were scored between the *HoxA* cluster and its regulatory regions in leg buds than that in wing buds (black rectangle). Expression levels were normalized to *Gapdh* and are shown as fold change relative to forelimb buds at either E10.5, or HH20-21. Error bars indicate standard deviation of either three (chick), two (E10.5) or four (E10.75) biological replicates. **P<0.01, *P<0.05, NS, P>0.05, Student’s *t*-test.

**S7 Fig (related to Fig 7).**
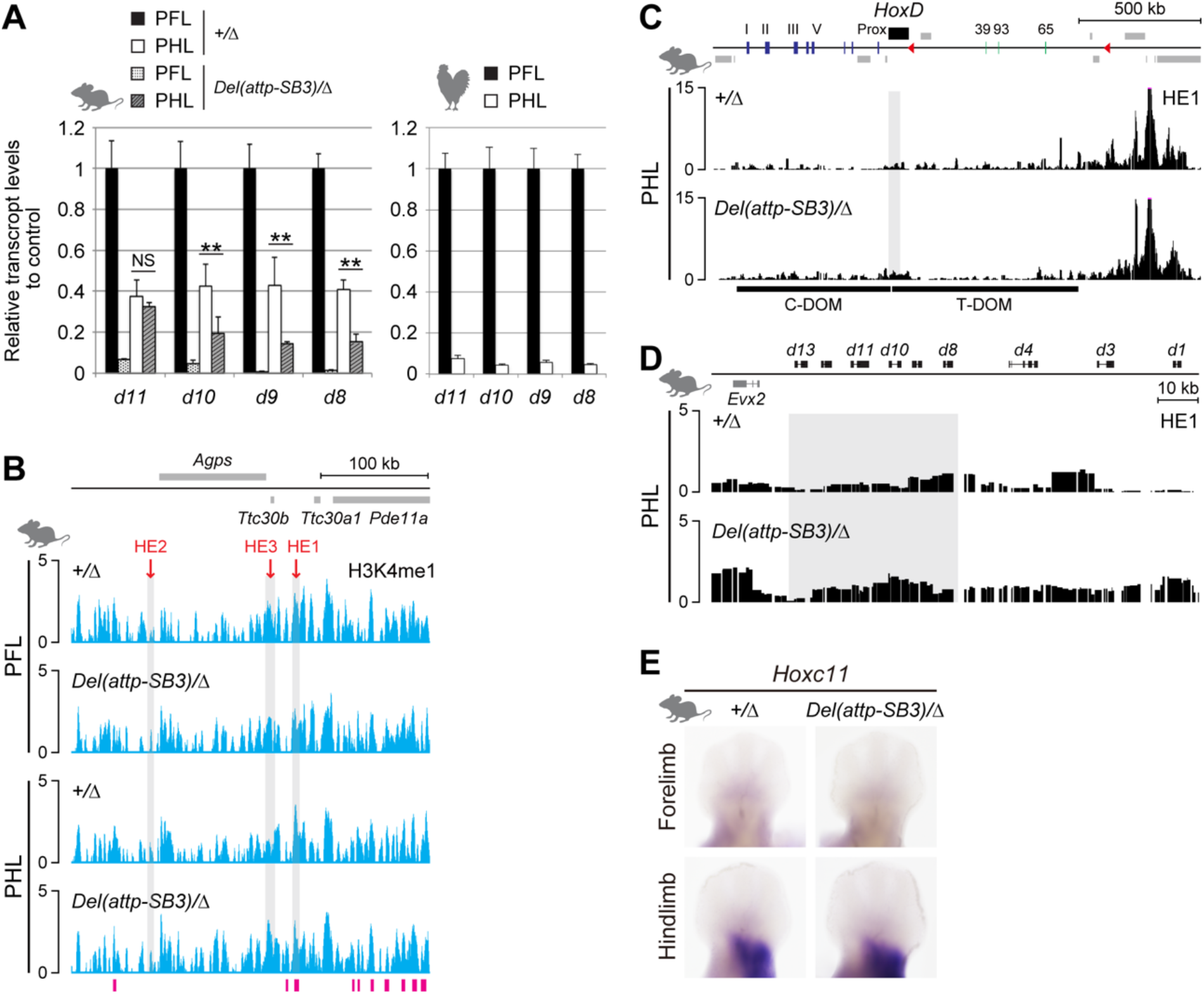
A T-DOM deletion induces interactions between HE1 and *Hoxd* genes. (A) Relative expression levels for each *Hoxd* gene in mouse and chick proximal fore-and hindlimbs. Expression levels in mouse and chick proximal fore-or hindlimb buds were normalized to *mGapdh* and *chGapdh*, respectively and are shown as fold change relative to mouse control or chick proximal forelimbs at E12.5 or HH28. Error bars indicate standard deviation of three (control), two (mutant) or three (chick) biological replicates. **P<0.01, NS, P>0.05, Student’s *t*-test. (B) H3K4me1 profiles obtained from proximal fore-and hindlimb buds of either control or *Del(attp-SB3)/∆* mutant embryos at E12.5. The putative HE1 enhancer was covered by H3K4me1 marks and merged with a predicted enhancer region. (C, D) 4C-seq profiles with HE1 as a viewpoint, by using either control, or *Del(attp-SB3)/∆* mutant proximal hindlimb buds at E12.5. The region from *Hoxd8* to *Hoxd13* is shaded (C). (D) Enlargement of (C) focusing on the *HoxD* cluster. Within the *Hoxd8* to *Hoxd13* interval, more contacts between HE1 and *Hoxd* promoters were detected in the mutant material where *Hoxd* expression remained as compared to the control situation where *Hoxd* expression was severely down-regulated. (E) *Hoxc11* expression from control and *Del(attp-SB3)/∆* mutant at E12.5. (left) Expression of *Hoxc11* in proximal hindlimb buds partly overlapped with that of *Hoxd11*. The deletion of T-DOM did not affect *Hoxc11* expression. Enrichment (*Y-*axis) of ChIP is shown at the log_2_ ratio of the normalized number of reads between ChIP and input samples. FL, forelimb; HL, hindlimb.

**S1 Table.**
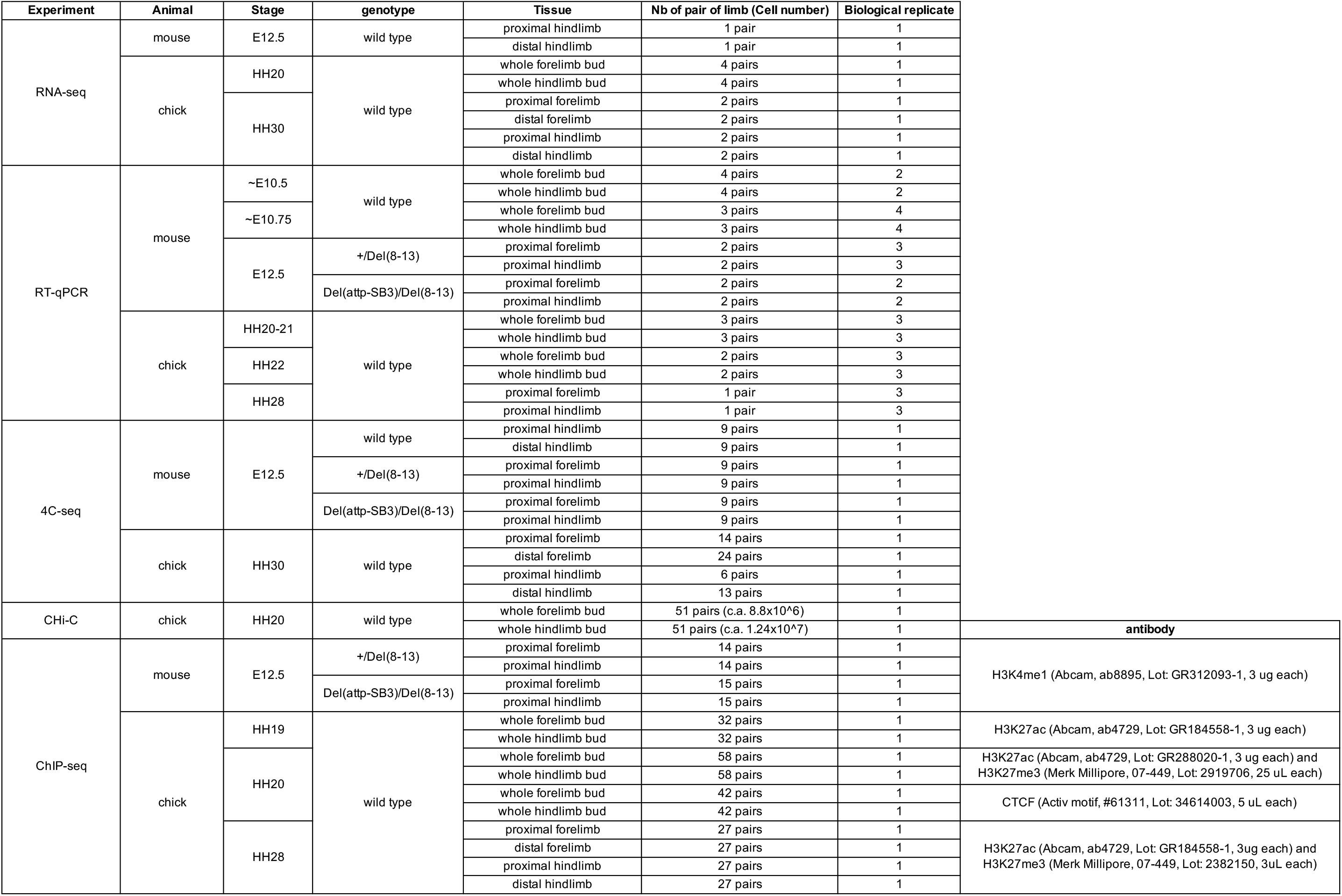
Information about samples.

**S2 Table.**
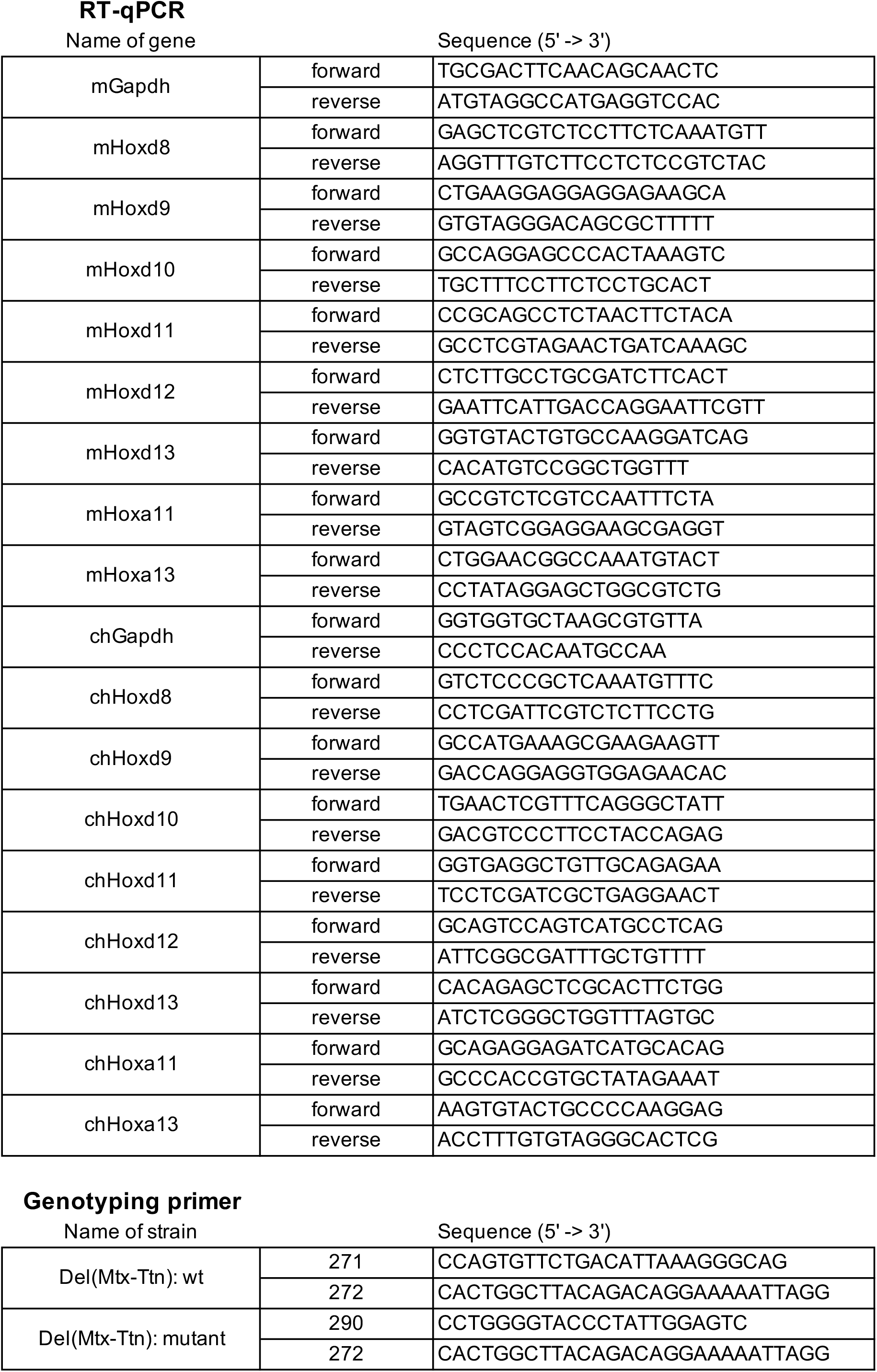
DNA sequences of primers used for both RT-qPCR analyses and genotyping.

**S3 Table.**
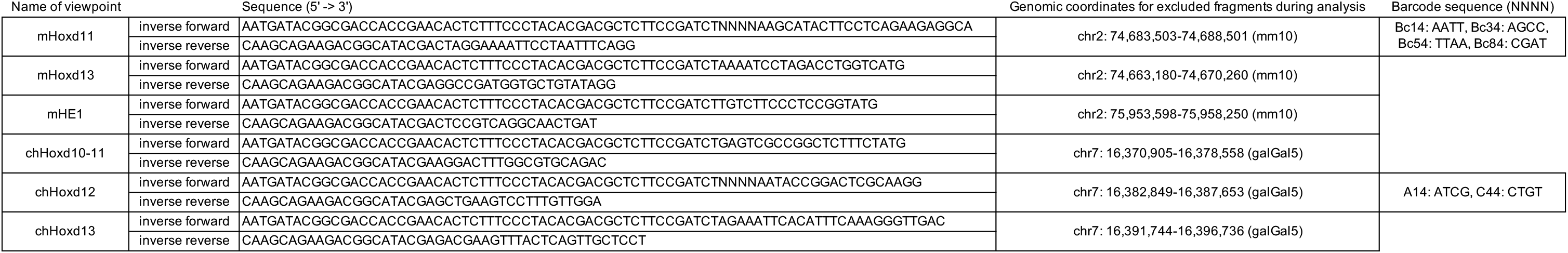
DNA sequences of primers used for 4C analyses. Custom barcodes (4bp shown by NNNN) were introduced in between the Illumina adapter sequence and the specific viewpoint sequences to do multiplexing using different samples with the same viewpoint.

**S4 Table.**
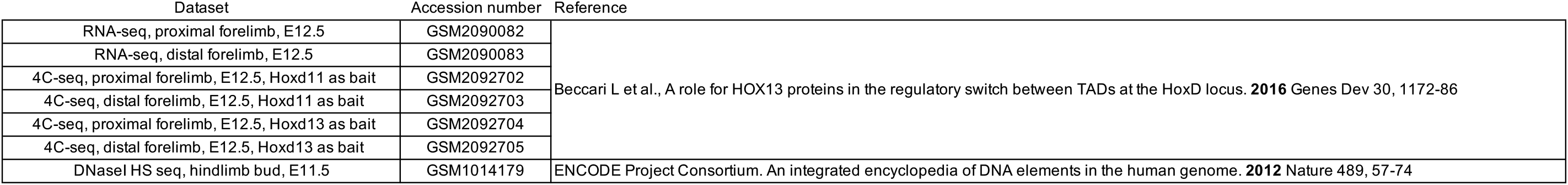
Public datasets used in this research.

